# Developmental *Gfi1* Dynamics Define Hematopoietic Emergence and Adult Hematopoietic Stem Cell Potency

**DOI:** 10.64898/2026.07.28.741133

**Authors:** Tomohiro Yabushita, Yosuke Tanaka, Takako Ideue, Kanako Wakahashi, Mariko Tsuruda, Saori Morino-Koga, Tomomasa Yokomizo, Taichi Noda, Akira Nishiyama, Daisuke Kurotaki, Terumasa Umemoto, Satoshi Yamazaki, Hitoshi Takizawa, Tomohiko Tamura, Minetaro Ogawa, Toshio Suda

## Abstract

*Gfi1* regulates endothelial-to-hematopoietic transition (EHT) and hematopoietic stem cell (HSC) maintenance; however, its expression dynamics remain unclear. We generated a non-disruptive *Gfi1*-T2A-mScarlet reporter to track *Gfi1* expression. At E10.5, reporter selectively marked EHT in the dorsal aorta, umbilical artery, and vitelline artery. At E9.5, mScarlet-positive endothelial cells were present in the umbilical and vitelline arteries but were scarce in the dorsal aorta, indicating vascular-bed-specific differences in hemogenic activation. *Gfi1* was broadly expressed in fetal liver HSCs, while higher expression levels identified HSCs with enhanced multilineage reconstitution and preferential T-cell output. Although *Gfi1* expression declined during fetal-to-adult maturation, a subset of adult bone marrow HSCs retained expression and exhibited superior repopulating and self-renewal capacity. Bulk and single-cell transcriptomic analyses linked this subset to dormant and fetal-associated programs, including imprinted genes. Reduced chromatin accessibility at a conserved *Gfi1* +26.5kb putative enhancer correlated with developmental *Gfi1* downregulation. Thus, *Gfi1* dynamics define EHT onset and functionally distinct HSC stemness.

## Introduction

Hematopoietic stem cells (HSCs) reside at the apex of the hematopoietic hierarchy, sustaining blood production through self-renewal and multilineage differentiation. HSCs are functionally heterogeneous and can be prospectively enriched by markers reflecting distinct transcriptional and signaling states^1–5^. *Growth factor independent 1* (*Gfi1*), a transcriptional repressor, is a key regulator of HSC function and was first identified in an interleukin-2-dependent T-cell line in the 1990s^6^.

During embryogenesis, definitive HSCs arise at embryonic day 10.5 (E10.5) from hemogenic endothelium (HE) located within the aorta-gonad-mesonephros (AGM) region. This process, known as the endothelial-to-hematopoietic transition (EHT), transforms endothelial cells into hematopoietic progenitors. *Runx1*, a central transcription factor in definitive hematopoiesis, induces the expression of *Gfi1* and *Gfi1b*, which promotes the loss of endothelial identity and initiates the hematopoietic program^7^. Consistent with this function, simultaneous loss of *Gfi1* and *Gfi1b* leads to sustained endothelial gene expression and prevents hematopoietic cells from budding and detaching from the endothelium, resulting in defective EHT. Previous studies investigating *Gfi1* expression during HSC development have primarily utilized *Gfi1*-or *Gfi1b*-reporter mice^7,8^. These reporter models were generated using fluorescent knock-in strategies that disrupt one endogenous allele of *Gfi1* and/or *Gfi1b*, which limits the ability to accurately visualize physiological *Gfi1* expression levels and dynamics. Furthermore, most analyses have focused on the AGM^8^, while the temporal and spatial patterns of *Gfi1* expression in other major tissues and vascular hematopoietic sites, including the yolk sac (YS), umbilical artery (UA), and vitelline artery (VA), remain poorly characterized.

Gfi1 is also important in adult HSC maintenance. Generally, transcription factors in HSCs do not act as simple binary switches, but rather exert quantitatively regulated effects depending on expression level^2,9–11^. This concept of dose-dependent regulation has been illustrated by factors governing HSC homeostasis. For instance, both insufficient and excessive expression of key regulators such as Evi1 and Runx1 perturb HSC emergence, underscoring the presence of an optimal expression window^2,10^. In this context, current Gfi1 reporter mouse models, which suffer from haploinsufficiency, are inherently limited in their ability to accurately resolve physiological expression levels, fine-scale gradients, or cellular heterogeneity *in vivo*. The regulatory basis of developmental changes in *Gfi1* expression also remains incompletely understood.

To overcome these limitations, we generated a *Gfi1*-T2A-mScarlet reporter mouse in which a self-cleaving T2A peptide enables fluorescent tagging of *Gfi1* expression without perturbing endogenous Gfi1 function. Using this model, we mapped the spatiotemporal dynamics of *Gfi1* across hematopoietic development and integrated functional, transcriptomic, and chromatin accessibility analyses to define how developmental *Gfi1* dynamics relate to HSC heterogeneity. These results demonstrated that *Gfi1* selectively marks cells at the EHT stage, identifies functionally distinct fetal and adult HSC subsets, and is associated with developmental remodeling of a conserved *cis*-regulatory element at the *Gfi1* locus.

## Results

### A physiological Gfi1 reporter selectively captures arterial EHT

To visualize endogenous *Gfi1* expression, we generated a *Gfi1*-T2A-mScarlet knock-in reporter mouse by inserting the reporter cassette immediately upstream of the endogenous *Gfi1* stop codon **(Figure 1A and Figure S1A)**. Correct targeting and normal steady-state peripheral blood hematopoiesis were confirmed **(Figure S1B-C)**. We first examined *Gfi1* expression at the onset of definitive hematopoiesis.

**Figure 1.**
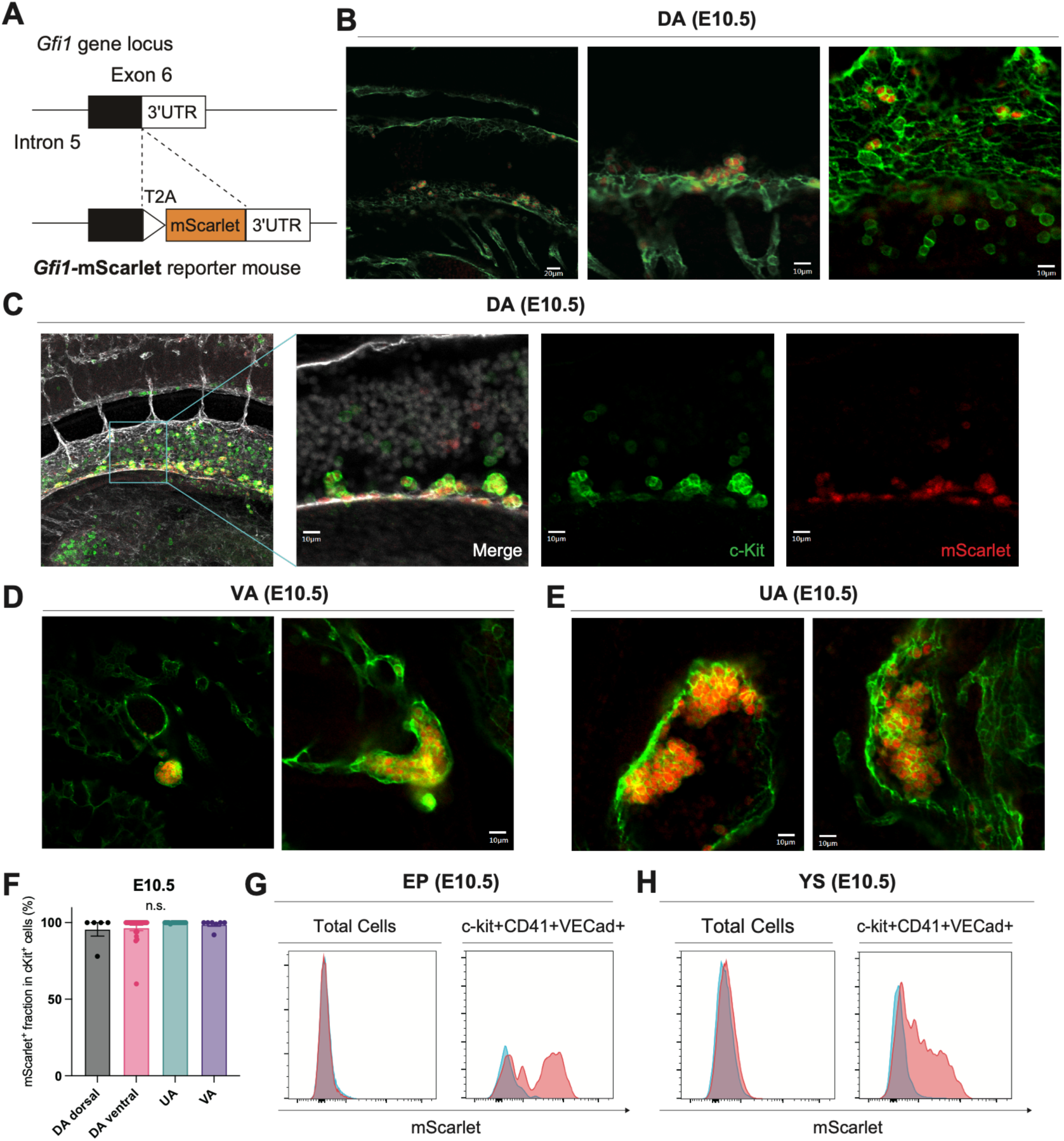
*Gfi1*-mScarlet reporter mouse visualizes the emergence of HSCs in the AGM regions. **(A)** Schematic representation of the *Gfi1*-T2A-mScarlet reporter allele, in which the T2A-mScarlet cassette was inserted immediately upstream of the stop codon in exon 6 of Gfi1. **(B)** Confocal images of the dorsal aorta (AGM region) in an E10.5 *Gfi1*-T2A-mScarlet embryo. Left, overview; middle, single optical slice; right, representative Z-stack image. **(C)** Confocal images of the dorsal aorta showing c-Kit (green) and mScarlet (red). Left, low-magnification maximum intensity projection (MIP); right three panels, high-magnification single optical slices (merge, c-Kit, and mScarlet). **(D-E)** Confocal views of the vitelline and umbilical arteries. Left, cross-sectional view; right, longitudinal view along the vascular axis. **(F)** Quantification of mScarlet-positive cells among morphologically defined IAHCs in the dorsal aorta (DA) and hematopoietic clusters in vitelline artery (VA) and umbilical artery (UA). **(G-H)** Flow cytometric analysis of E10.5 *Gfi1*-T2A-mScarlet embryos. (E) Embryo proper (EP); (F) yolk sac (YS). Left, total cells; right, c-Kit⁺CD41⁺VE-Cad⁺ population. Red, *Gfi1*-mScarlet +/WT; blue, WT controls.

At E10.5, definitive hematopoiesis emerges from the dorsal aorta (DA) within the aorta-gonad-mesonephros (AGM) region. Confocal imaging of *Gfi1*-mScarlet heterozygous embryos revealed strong mScarlet fluorescence in intra-aortic hematopoietic clusters (IAHCs) protruding from the DA endothelium, whereas wild-type embryos showed no detectable signal **(Figure 1B-C; Figure S2A-B)**. These c-Kit⁺ IAHCs were located predominantly on the ventral side of the DA, with occasional smaller buds or clusters observed dorsally, and reached up to ∼10 μm in diameter, consistent with pre-HSCs and progenitors generated through EHT **(Figure 1C)**^12^. Along the DA, mScarlet⁺ endothelial cells showed a discontinuous, patchy pattern, particularly in the middle ventral region, with c-Kit⁺ IAHCs located at discrete sites within these mScarlet⁺ endothelial areas. In contrast, neither fetal liver nor non-hematopoietic tissues in the embryo proper exhibited detectable mScarlet fluorescence, indicating that Gfi1 reporter activity at E10.5 was highly restricted to IAHCs and associated endothelial cells **(Figure S2C-D)**.

Reporter activity was also detected in hematopoietic clusters in the vitelline artery (VA) and umbilical artery (UA). These clusters were larger than those in the DA, reaching up to ∼50μm in diameter **(Figure 1D-E, Figure S2E-F)**. mScarlet⁺ cells bulged from the luminal surface of the endothelium, whereas adjacent cluster-free endothelial cells were mScarlet-negative. Hematopoietic clusters in the DA, VA, and UA were detected with near-complete sensitivity by the *Gfi1*-mScarlet reporter **(Figure 1F)**, indicating that virtually all morphologically identifiable arterial cluster-forming cells expressed *Gfi1* at this stage. Confocal evaluation of mScarlet in the yolk sac (YS) was limited by background fluorescence associated with anti-RFP staining. Flow cytometric analysis revealed that mScarlet-positive cells were extremely rare (<0.1%) among total cells in the embryo proper **(Figure 1G)**. However, when gating on VE-cadherin⁺CD41⁺c-Kit⁺ cells undergoing EHT, approximately half were mScarlet⁺ **(Figure 1G)**, supporting the selective association of *Gfi1* expression with ongoing EHT. A similar pattern was observed in the YS: although mScarlet⁺ cells constituted <0.1% of total cells, ∼30-40% of EHT-ongoing cells were mScarlet-positive **(Figure 1H)**.

Because the reporter expression in the YS could not be reliably evaluated by confocal microscopy, endogenous Gfi1 protein was further assessed using an anti-Gfi1 antibody. Gfi1 immunostaining confirmed expression in subsets of IAHCs and DA endothelial cells **(Figure S3A-B)**, as well as in portions of c-Kit⁺ clusters in the VA and UA **(Figure S3C)**. In the YS, anti-Gfi1 staining exhibited minimal background and revealed heterogeneous Gfi1 expression among c-Kit⁺ clusters, with both strongly positive and negative clusters **(Figure S4A-B)**. Collectively, these findings indicate that at E10.5, Gfi1 expression is tightly associated with ongoing EHT, marking IAHCs and associated endothelial cells in the DA as well as hematopoietic clusters in the VA and UA, while showing more heterogeneous expression in the YS. Thus, the *Gfi1*-mScarlet reporter provides a robust *in vivo* marker of EHT in arterial hematopoietic sites.

### Gfi1 Marks Early Hemogenic Activation preferentially in the UA and VA

The precise timing of hemogenic specification in different embryonic vascular beds is still under debate. Although several studies have reported that EHT in the UA and VA peaks around E10.5 concurrently with the DA, the temporal relationship among these regions remains unresolved. To address this issue, we focused on an earlier developmental stage (E9.5) and investigated the onset of *Gfi1* expression in UA and VA endothelium prior to HSC emergence in the DA at E10.5.

Confocal imaging of *Gfi1*-mScarlet embryos revealed reproducible mScarlet-positive cells embedded within the CD31⁺ endothelial lining of both UA and VA at E9.5, before robust hematopoietic cluster formation in the DA at E10.5 **(Figure 2A-B)**. These mScarlet-positive endothelial cells were sparsely integrated within the CD31⁺ vascular lining of the UA and VA, without overt protrusive morphology or cluster formation, suggesting that Gfi1 upregulation precedes overt EHT morphology in these vessels. In contrast, reporter activity was minimal in the DA at the same stage **(Figure 2C)**, consistent with previous work showing that hematopoietic cells appear earlier in the VA and UA than in the DA^13^. Consistent findings were obtained by immunostaining wild-type E9.5 embryos: endogenous Gfi1 protein was detected as scattered or short, linear stretches of positive endothelial cells in UA and VA, whereas DA staining was minimal or undetectable **(Figure S5A)**, and isotype controls remained negative **(Figure S5B)**. These results strongly suggest that Gfi1 upregulation initiates earlier in the VA and UA than in the DA.

**Figure 2.**
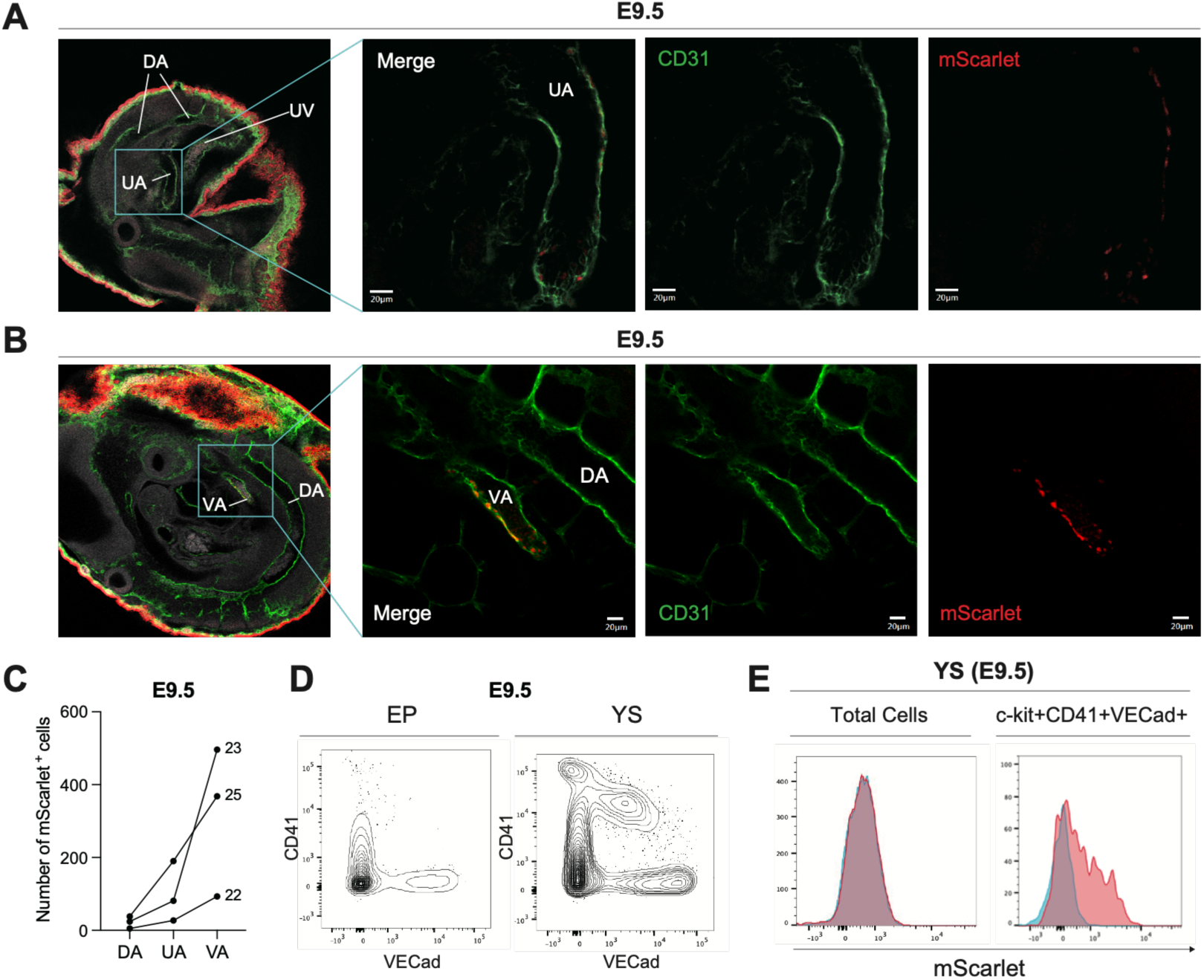
*Gfi1*-mScarlet reporter marks a subset of endothelial cells in the umbilical and vitelline arteries at E9.5. **(A-B)** Confocal images of E9.5 *Gfi1*-T2A-mScarlet embryos showing endothelial structures of the umbilical artery (UA) (A) and vitelline artery (VA) (B). CD31 (green) marks endothelial cells and mScarlet (red) indicates reporter expression. DA, dorsal aorta. **(C)** Number of mScarlet^+^ endothelial cells within the vascular endothelium of the umbilical artery (UA), vitelline artery (VA), and dorsal aorta (DA) in E9.5 embryos (n = 3). mScarlet^+^ endothelial cells were defined as mScarlet^+^cells integrated within the CD31^+^vascular lining. Data points from the same embryo are connected by lines, and the somite number for each embryo is indicated on the right (22, 23, and 25). **(D)** Flow-cytometric analysis of embryo proper (EP) and yolk sac (YS) from E9.5 Gfi1-T2A-mScarlet embryos. Each plot shows VE-cadherin (x-axis) versus CD41 (y-axis). **(E)** Flow-cytometric profiles of E9.5 embryo Yolk sac. Left, total cells; right, c-Kit⁺ CD41⁺ VE-cadherin⁺ population. Red, *Gfi1*-mScarlet *⁺/WT*; blue, WT controls.

To complement the imaging data, flow cytometry was performed to assess Gfi1 expression within the YS at E9.5. Whereas the embryo proper contained few if any VE-cadherin⁺CD41⁺c-Kit⁺ cells, the YS harbored a clear population of such EHT-stage cells, approximately one third of which were mScarlet-positive **(Figure 2D-E)**. Consistently, immunostaining of wild-type E9.5 embryos showed heterogeneous *Gfi1* expression within c-Kit⁺ clusters in the YS **(Figure S6A-B)**, further supporting early Gfi1 activation at this stage. Collectively, these findings demonstrate that although Gfi1-positive cells were extremely rare at E9.5, their emergence was spatially restricted to UA, VA, and YS hematopoietic clusters, and precedes detectable EHT in the DA. Although Gfi1 has frequently been linked to AGM-derived definitive hematopoiesis, our data reveal that *Gfi1* marks EHT cells independently of the hematopoietic wave or anatomical sites. The *Gfi1*-mScarlet reporter therefore provides a powerful *in vivo* readout of ongoing hemogenic specification. Moreover, these findings suggest that the hemogenic specification occurs earlier in the UA and VA before the DA enters its cluster-forming phase.

### *Gfi1* is highly expressed in fetal liver HSCs, and higher expression marks enhanced stemness and T-cell-biased output

By E12.5, HSCs colonize FL and undergo rapid expansion. To define *Gfi1* expression during this phase, FL cells from *Gfi1* reporter mice were analyzed by flow cytometry at E14.5, when HSCs can be reliably identified using adult-equivalent surface markers. At this stage, *Gfi1* expression was highest in long-term HSCs (LT-HSCs) and progressively decreased in short-term HSCs (ST-HSCs), multipotent progenitors (MPPs), myeloid progenitors (MyPs), and mature hematopoietic cells (Lin⁺) **(Figure 3A-B)**. This graded pattern supports a close association between high *Gfi1* expression and fetal HSC identity. At later fetal stages (E17.5 and E19.5), *Gfi1* expression increased in myeloid progenitors and differentiated cells while remaining high in LT-HSCs **(Figure S7A-C)**.

**Figure 3.**
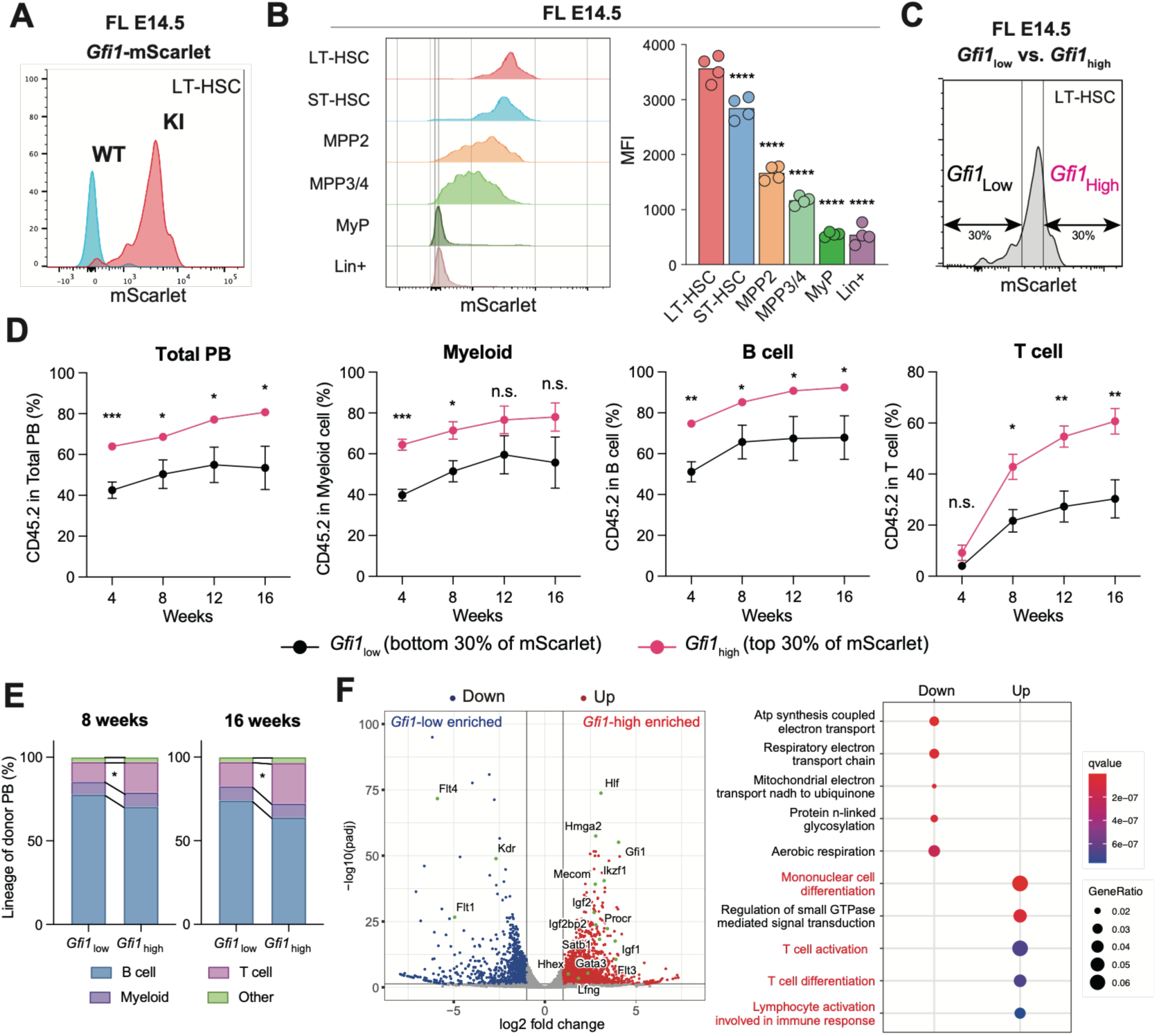
Elevated *Gfi1* expression in fetal liver HSCs associates with increased peripheral output, especially with a T-cell-lymphoid bias. **(A)** Representative histogram of mScarlet fluorescence in E14.5 fetal liver (FL) LT-HSCs. Red, *Gfi1*-mScarlet ⁺/WT; blue, WT controls. **(B)** Histograms (left) and quantification of mScarlet mean fluorescence intensity (MFI) (right) across indicated FL populations: LT-HSC, ST-HSC, MPP2, MPP3/4, myeloid progenitors (MyP), and Lin⁺ cells. **(C)** Representative flow cytometry plots defining *Gfi1*-high and *Gfi1*-low E14.5 FL HSCs as the top and bottom 30% of mScarlet fluorescence, respectively. **(D)** Competitive transplantation of FL *Gfi1*-high and *Gfi1*-low HSCs into Ly5.1 recipient mice. Donor-derived peripheral blood chimerism at 4, 8, 12, and 16 weeks post-transplantation showing total, myeloid, B-, and T-cell chimerism (n = 5). **(E)** Lineage distribution of donor-derived cells in peripheral blood at 8 and 16 weeks post-transplantation. **(F)** Volcano plot showing differentially expressed genes (DEGs) between *Gfi1*-high and *Gfi1*-low FL HSCs (n = 2; adjusted *P* < 0.05 and an absolute log₂ fold change > 2) (left). Gene Ontology (GO) enrichment analysis of biological processes enriched among the DEGs (right).

Although *Gfi1* expression was uniformly high in FL LT-HSCs relative to downstream progenitors, measurable heterogeneity persisted within the LT-HSC compartment. To assess the functional significance of this variation, E14.5 FL LT-HSCs were stratified by mScarlet intensity, with the top and bottom 30% defined as *Gfi1*-high and *Gfi1*-low, respectively, and subjected to competitive BM transplantation. *Gfi1*-high HSCs consistently showed superior multilineage reconstitution from 4 to 16 weeks post-transplantation, with significantly higher donor chimerism in both B and T cells **(Figure 3C, Figure S8A)**. The proportion of T cells among donor-derived CD45.2⁺ cells also increased after 8 weeks post-transplantation **(Figure 3D)**. At 16 weeks, donor-derived LT-HSCs were recovered at similar frequencies in recipients of both groups, whereas donor-derived MPPs were significantly more abundant in recipients of *Gfi1*-high HSCs **(Figure S8B)**. The reconstitution advantage of *Gfi1*-high HSCs persisted in secondary transplantation, with recipients showing significantly higher donor chimerism at both 2 and 4 months, most notably in the T-cell compartment **(Figure S8B-C)**.

To investigate whether this functional heterogeneity is associated with transcriptional differences, we performed RNA sequencing on E14.5 fetal liver HSCs stratified by Gfi1 expression **(Figure S9A-B)**. *Gfi1*-high fetal HSCs preferentially expressed fetal liver HSC-associated genes (*Hmga2* and *Hlf*), imprinted genes (*Meg3*, *Mirg*), and factors involved in stem cell maintenance and dormancy (*Mecom*, *Mllt3*), whereas *Gfi1*-low HSCs were enriched for endothelial-associated transcripts **(Figure 3F. Figure S9C)**. Gene ontology analysis further showed that *Gfi1*-high HSCs were enriched for lymphoid-associated programs, including mononuclear cell differentiation, T-cell activation, and T-cell differentiation, consistent with preferential T-cell reconstitution. In contrast, oxidative phosphorylation and aerobic respiration pathways were overrepresented among genes preferentially expressed in *Gfi1*-low HSCs, suggesting reduced stemness or increased metabolic activation **(Figure 3F, Figure S9D-E)**. Together, these results indicate that elevated *Gfi1* expression within fetal liver HSCs is associated with transcriptional programs linked to stemness and preferential T-cell-lymphoid output.

### *Gfi1* Expression Declines During HSC Development and identifies a Functionally Distinct Adult HSC Subset

At birth, hematopoiesis shifts from FL to BM, accompanied by a gradual transition in HSC intrinsic properties^14^. Comparative analysis of mScarlet reporter fluorescence in FL LT-HSCs and adult BM LT-HSCs demonstrated a significant overall reduction in *Gfi1* expression in adult HSCs. Although most adult LT-HSCs showed mScarlet fluorescence near wild-type background levels, approximately 30% retained intermediate but clearly detectable expression, which remained substantially lower than those of FL LT-HSCs **(Figure 4A)**. Unlike E14.5 FL-HSC, which expressed more Gfi1 than downstream progenitors, adult BM LT-HSCs showed lower average *Gfi1* expression than early progenitor and mature cell populations **(Figure 4B)**, reflecting strong Gfi1 expression in granulocyte-macrophage progenitors (GMPs), neutrophils, eosinophils, and Ly6C⁺ monocytes within adult BM **(Figure S10A-D)**. To assess whether residual Gfi1 expression defines a functionally distinct adult HSC subset, adult LT-HSCs were sorted by mScarlet intensity into *Gfi1*-high (top 30%) and *Gfi1*-low (bottom 30%) fractions and subjected to competitive transplantation. *Gfi1*-high adult BM HSCs showed significantly enhanced long-term multilineage reconstitution compared with *Gfi1*-low HSCs, maintaining higher donor-derived blood chimerism across all lineages through 16 weeks post-transplantation **(Figure 4C)**. BM analysis further showed that recipients of *Gfi1*-high HSCs consistently exhibited greater donor contributions in LT-HSCs, ST-HSCs, and MPPs **(Figure 4D)**. Moreover, at 16 weeks, donor-derived LT-HSCs recovered from recipients of *Gfi1*-high HSCs contained a significantly higher fraction of mScarlet-positive cells than those from recipients of *Gfi1*-low HSCs, suggesting better preservation of a primitive HSC state after transplantation **(Figure 4E)**. This advantage persisted after secondary transplantation, consistent with superior self-renewal capacity of *Gfi1*-high adult HSCs **(Figure 4F, Figure S11A-B)**. Together, these findings indicate that, despite the overall reduction of *Gfi1* expression in adult HSCs, the relatively *Gfi1*-high subset drives sustained long-term hematopoietic reconstitution.

**Figure 4.**
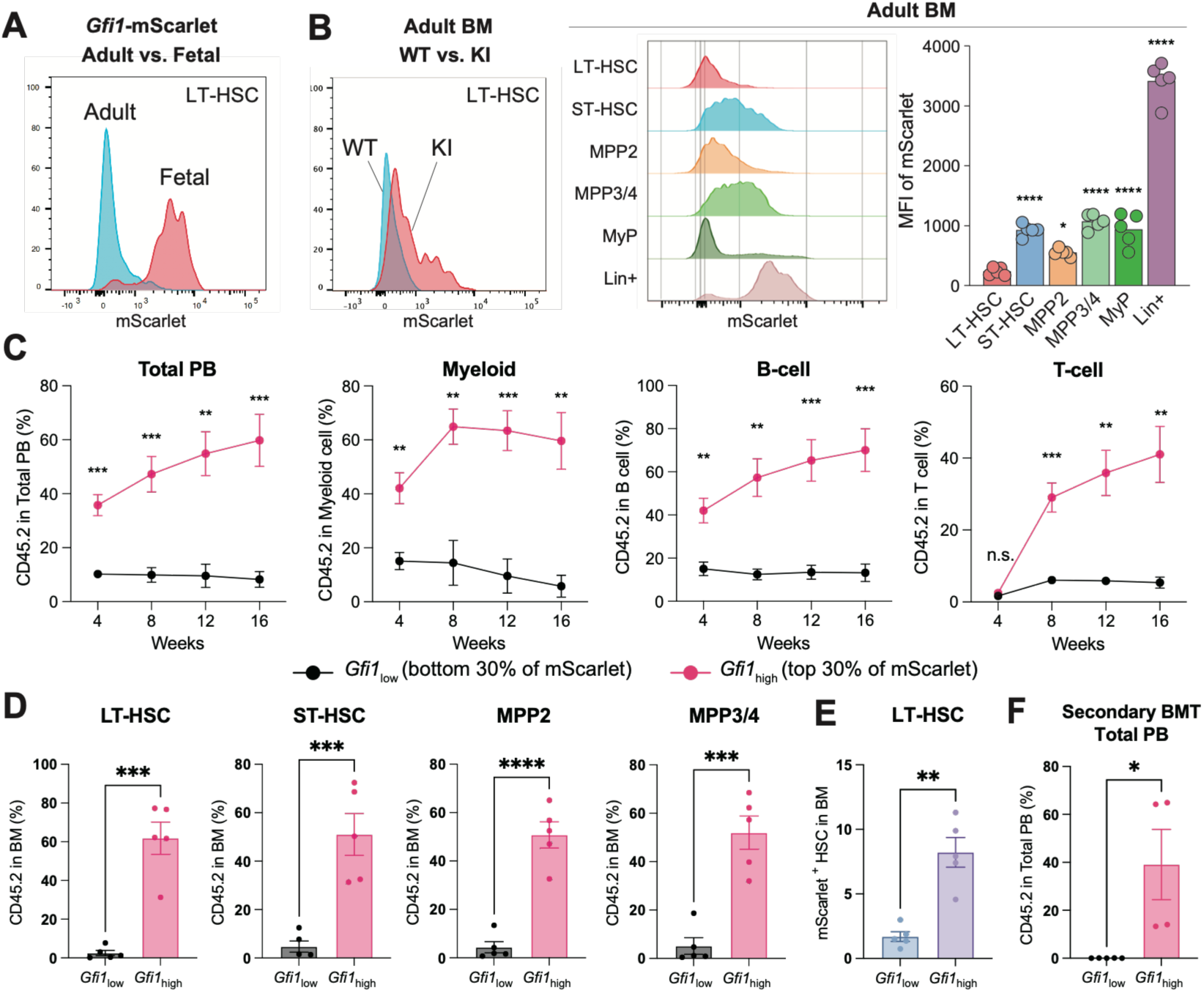
*Gfi1* is broadly downregulated in adult HSCs, whereas its residual expression marks HSCs with robust repopulating capacity. **(A)** Direct comparison of mScarlet fluorescence between FL LT-HSCs and adult BM LT-HSCs from *Gfi1*-mScarlet *+/*WT mice. **(B)** Representative histogram of mScarlet fluorescence in adult bone marrow (BM) LT-HSCs. Red, *Gfi1*-mScarlet*+/*WT; blue, WT controls (left). Histograms (middle) and comparison of mScarlet MFI (right) among adult BM populations: LT-HSC, ST-HSC, MPP2, MPP3/4, MyP, and Lin⁺ cells. **(C)** Peripheral blood chimerism following competitive transplantation of adult BM *Gfi1*-high vs. *Gfi1*-low HSCs at 4-16 weeks, showing total, myeloid, B-, and T-cell contributions (n = 5). **(D)** Bone marrow chimerism at 16 weeks, measured in LT-HSC, ST-HSC, MPP2, and MPP3/4 subsets from the same recipients as in (C). **(E)** Frequency of mScarlet-positive cells among donor-derived BM HSCs at 4 months after transplantation of adult BM *Gfi1*-high or *Gfi1*-low HSCs. **(F)** Total donor-derived peripheral blood chimerism at 16 weeks after secondary transplantation of adult BM HSCs.

Bulk RNA sequencing revealed that adult *Gfi1*-high and *Gfi1*-low HSCs possess distinct transcriptional profiles **(Figure 5A–C)**. To associate these transcriptional differences with functional states, gene set enrichment analysis (GSEA) was conducted. *Gfi1*-high HSCs were significantly enriched for a LT-HSC signature^15^ and the dormant HSC signature (MolO signature^16^) **(Figure 5C)**. Consistent with these transcriptional features, EPCR (Procr) expression was increased in *Gfi1*-high HSCs, as confirmed by flow cytometry **(Figure 5D)**. Conversely, *Gfi1*-low HSCs were significantly enriched for the Primed HSC signature (NoMO signature^16^), which is associated with lineage priming and activation states **(Figure 5C)**. Notably, fetal liver HSC-associated gene sets^17^ were selectively enriched in *Gfi1*-high adult HSCs, with prominent upregulation of imprinting-related genes, including *H19*, *Igf2*, *Meg3*, and *Mirg* **(Figure 5E-F)**. Because imprinted gene programs have been associated with LT-HSCs similar to other somatic stem cell populations^18^, these findings suggest that *Gfi1*-high adult HSCs exhibit a developmentally associated transcriptional state characterized by fetal-associated and imprinting-related features.

**Figure 5.**
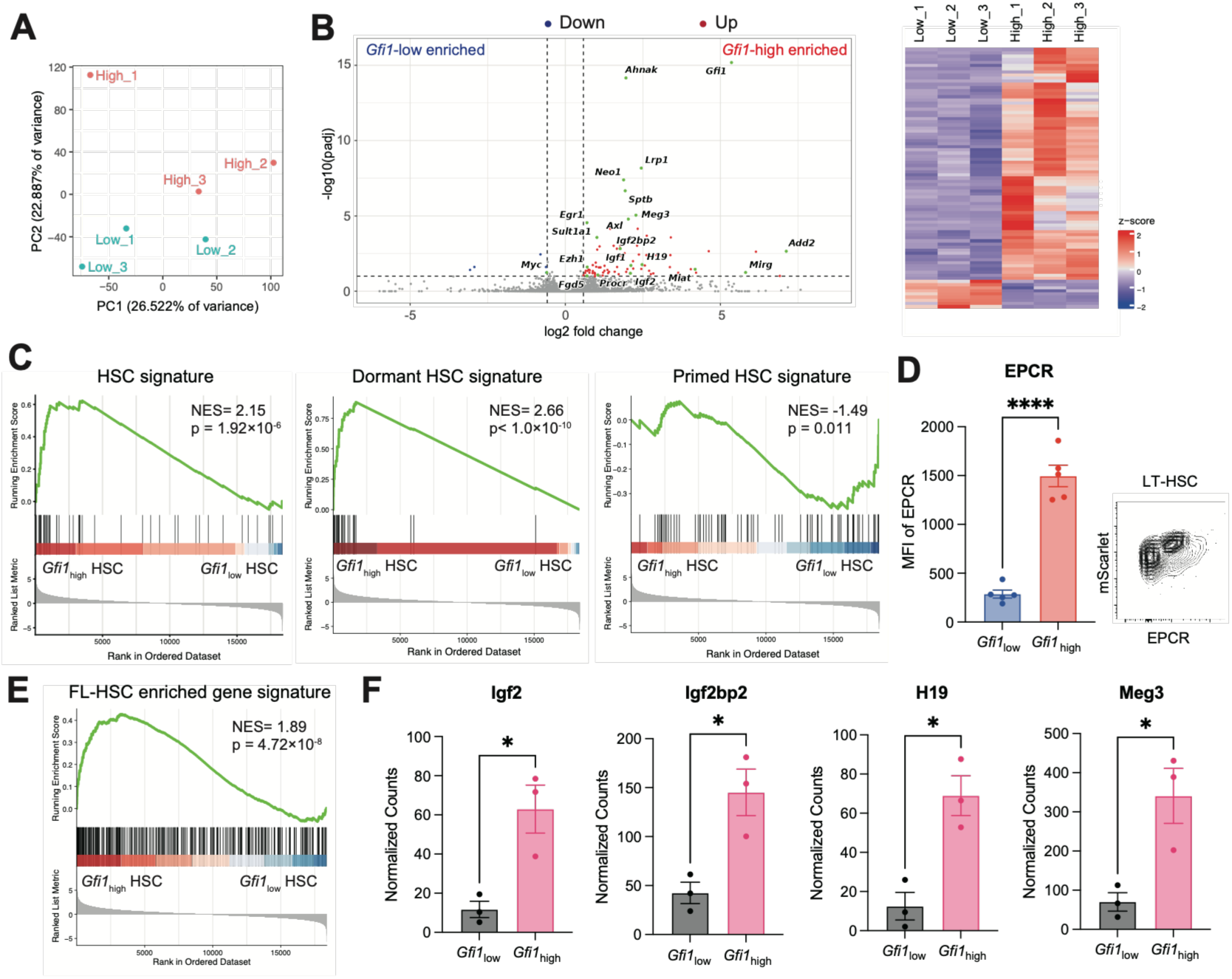
*Gfi1*-high adult HSCs exhibit dormant and fetal-associated molecular signatures. **(A)** Principal component analysis (PCA) of transcriptomic profiles from *Gfi1*-high and *Gfi1*-low adult BM HSCs (n = 3). **(B)** Volcano plot and hierarchical clustering heatmap showing differentially expressed genes (DEGs) between *Gfi1*-high and *Gfi1*-low adult BM HSCs (n = 3). DEGs were defined by an adjusted *P* value < 0.1 and an absolute log₂ fold change > 1.5. **(C)** Pre-ranked GSEA of RNA-seq data from *Gfi1*-high and *Gfi1*-low adult HSCs using HSC, dormant HSC, and primed HSC gene signatures^15,16^. **(D)** Comparison of EPCR mean fluorescence intensity (MFI) between *Gfi1*-high and *Gfi1*-low adult BM HSCs (left). Representative flow cytometry plots showing EPCR and mScarlet expression within the HSC population (right). **(E)** Preranked GSEA of RNA-seq data from *Gfi1*-high and *Gfi1*-low adult BM HSCs using FL HSC–enriched gene signatures^17^. **(F)** Normalized RNA-seq counts of representative FL HSC-enriched genes (*Igf2, Igf2bp2,* H19, and *Meg3*) in adult BM *Gfi1*-high and *Gfi1*-low HSCs (n = 3).

### *Gfi1* expression becomes progressively restricted to a dormant HSC subset during hematopoietic development

We integrated previously published scRNA-seq datasets^14^ of LT-HSCs isolated from fetal liver (E16.5), neonatal stages (P7 and P14), and adult bone marrow. UMAP visualization showed that HSCs did not form discrete clusters but instead were distributed along a developmentally ordered continuum spanning fetal, neonatal, and adult stages **(Figure 6A)**. Analysis of Gfi1 expression along the developmental axis revealed peak expression at the fetal stage (E16.5), with a gradual decline through the neonatal stages into adulthood **(Figure 6B)**. A subset of adult BM HSCs retained relatively high *Gfi1* expression, consistent with the *Gfi1*-high population identified in the reporter mouse. To further investigate this decline, we compared Gfi1 expression with that of key HSC self-renewal transcription factors (*Mecom*, *Gata2*, and *Hlf*). From E16.5 to adulthood, *Gfi1* expression decreased by ∼9.5-fold, whereas *Mecom* and *Hlf* expression declined by ∼2.2-fold and ∼1.7-fold, respectively, and Gata2 expression increased by ∼1.6-fold **(Figure 6B-C)**. Similarly, fetal HSC-associated and imprinting-related genes enriched in *Gfi1*-high HSCs, including *H19*, *Igf1*, *Igf2bp2*, and *Meg3*, were also markedly downregulated, indicating progressive attenuation of fetal-associated transcriptional programs toward adulthood **(Figure 6D)**^17,18^.

**Figure 6.**
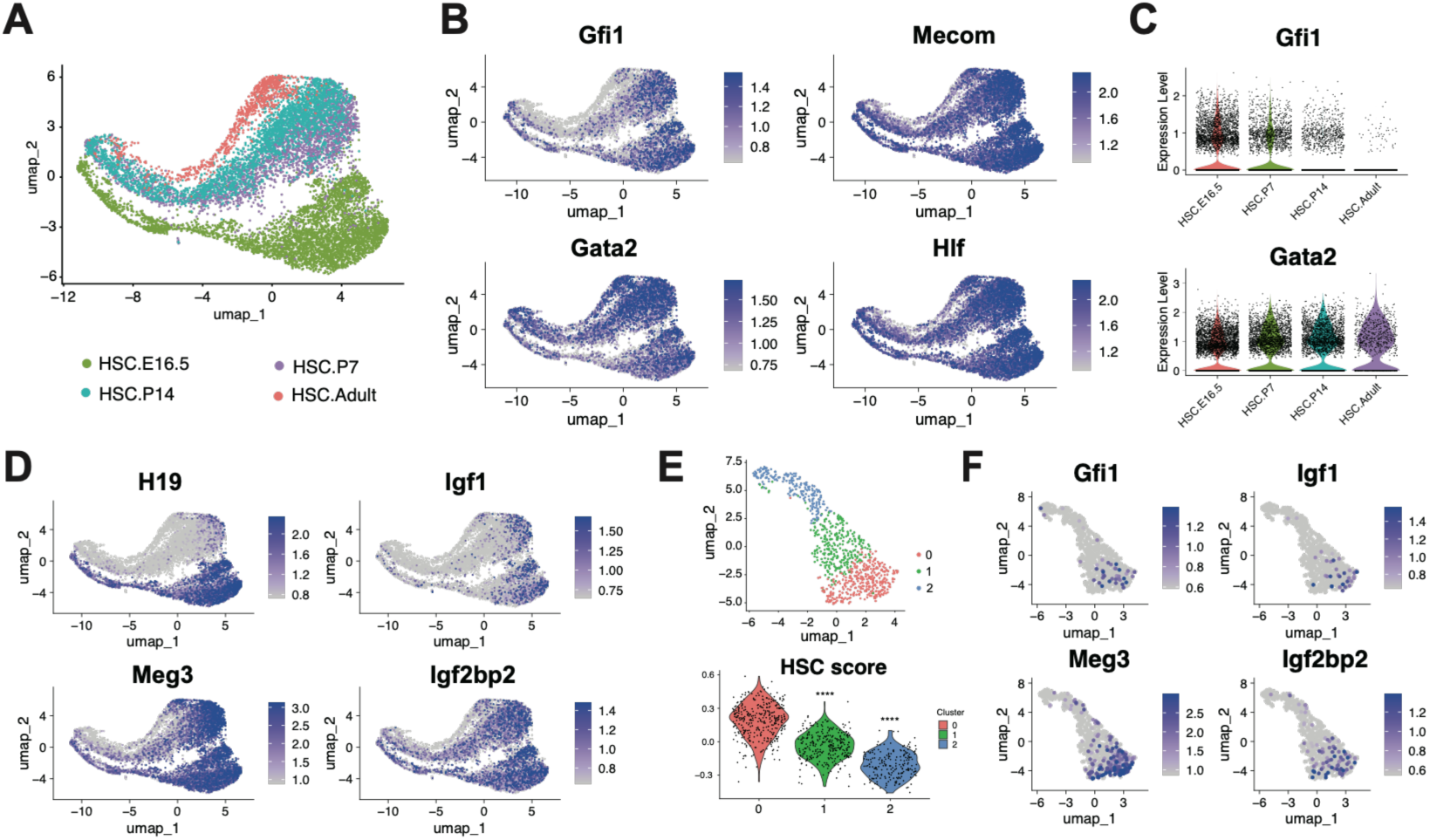
scRNA-seq shows high *Gfi1* expression in fetal HSCs that declines with age, leaving a potent Gfi1-positive subset in adults. **(A)** UMAP generated from an integrated analysis of a reanalyzed public scRNA-seq dataset^14^ containing fetal HSCs (E16.5), neonatal HSCs (P7, P14), and adult HSCs. **(B)** Expression of *Gfi1* and HSC-associated transcription factors (*Mecom, Gata2, Hlf*) visualized on the UMAP in (A). **(C)** Violin plots showing normalized expression of *Gfi1* (top) and *Gata2* (bottom). **(D)** UMAP feature plots of fetal liver–enriched genes (*H19, Igf1, Meg3, Igf2bp2*) showing preferential localization in Gfi1-high HSC populations. **(E)** Adult HSC subset extracted from (A): UMAP showing cluster identity (top) and violin plots showing the HSC dormancy score (MolO score^16^) across clusters (bottom). **(F)** Feature plots of *Gfi1, Igf1, Meg3,* and *Igf2bp2* in adult HSCs as displayed in (E).

To further clarify stage-specific heterogeneity, E16.5 and P7 HSCs were independently subclustered and evaluated using the dormant HSC signature (MolO signature^16^) **(Figure S12 A-D)**. At both stages, clusters with the highest HSC scores showed only modestly higher *Gfi1* expression (∼1.3-fold at E16.5 and ∼1.7-fold at P7), indicating that Gfi1 remained broadly expressed across fetal and neonatal HSCs **(FigureS12 A, C)**. In contrast, adult HSCs showed a distinct pattern, with *Gfi1* expression enriched 3.6-fold in the most dormant HSC cluster (p = 0.003) **(Figure 6E)**. This cluster exhibited significantly higher expression of established dormant HSC markers, including *Mllt3* (p = 6.8 × 10⁻³³), *Procr* (p = 8.9 × 10⁻¹⁰), *Ly6a* (p = 2.6 × 10⁻¹⁵), and *Vwf* (p = 3.8 × 10⁻⁵), as well as fetal HSC-associated genes identified by bulk RNA-seq, including imprinting-related genes such as *Meg3*, *H19*, and *Mirg*, together with *Igf1*, *Igf2bp2*, and *St8sia4* **(Figure 6F, Figure S12E)**.

### Chromatin accessibility at the *Gfi1* +26.5-kb putative enhancer decreases during fetal-to-adult HSC development

To investigate the regulatory basis of developmental *Gfi1* downregulation, we assessed chromatin accessibility in *Gfi1*-high and *Gfi1*-low HSCs isolated from E19.5 FL and adult BM. ATAC-seq profiles centered on genome-wide annotated transcription start sites (TSS) were comparable among the four groups; accessibility at the *Gapdh* TSS was also unchanged **(Figure 7A, Figure S13A-B)**. We next focused on two distal *cis*-regulatory elements **(Figure 7B)**: the -33.5 kb early hematopoietic enhancer^19^, active in AGM and FL reporter assays, and a conserved +26.5 kb region corresponding to a *GFI1* super-enhancer element in human acute erythroid leukemia cells^20^. Accessibility at the *Gfi1* TSS was significantly lower in adult *Gfi1*-low HSCs than in fetal *Gfi1*-high and *Gfi1*-low HSCs, consistent with their lower *Gfi1* expression **(Figure 7C-D)**. Accessibility at the +26.5 kb region was lowest in adult *Gfi1*-low HSCs and was significantly reduced compared with fetal *Gfi1*-high and *Gfi1*-low HSCs **(Figure 7C,E)**. This region was enriched for H3K27ac and H3K4me1 in primary fetal and adult HSPCs and occupied by *Gata2*, *Tal1*, and *Erg* in Hoxb8-FL HSPC cell lines^21^, supporting its designation as a putative enhancer **(Figure 7C)**. By contrast, accessibility at the -33.5 kb enhancer was not reduced during the fetal-to-adult transition **(Figure S13C-D)**, indicating region-specific remodeling of the *Gfi1* locus. To assess whether these changes extended to downstream components of the GFI1-GATA2-GFI1B regulatory circuit^22^, we examined the GFI1-bound *Gata2* −83 kb regulatory element and the *Gfi1b* +16 kb enhancer. Accessibility at both regions was inversely associated with *Gfi1* expression across the HSC populations **(Figure S13E-F)**. These findings identify reduced accessibility at the +26.5-kb putative enhancer during fetal-to-adult HSC development as a chromatin feature associated with *Gfi1* downregulation.

**Figure 7.**
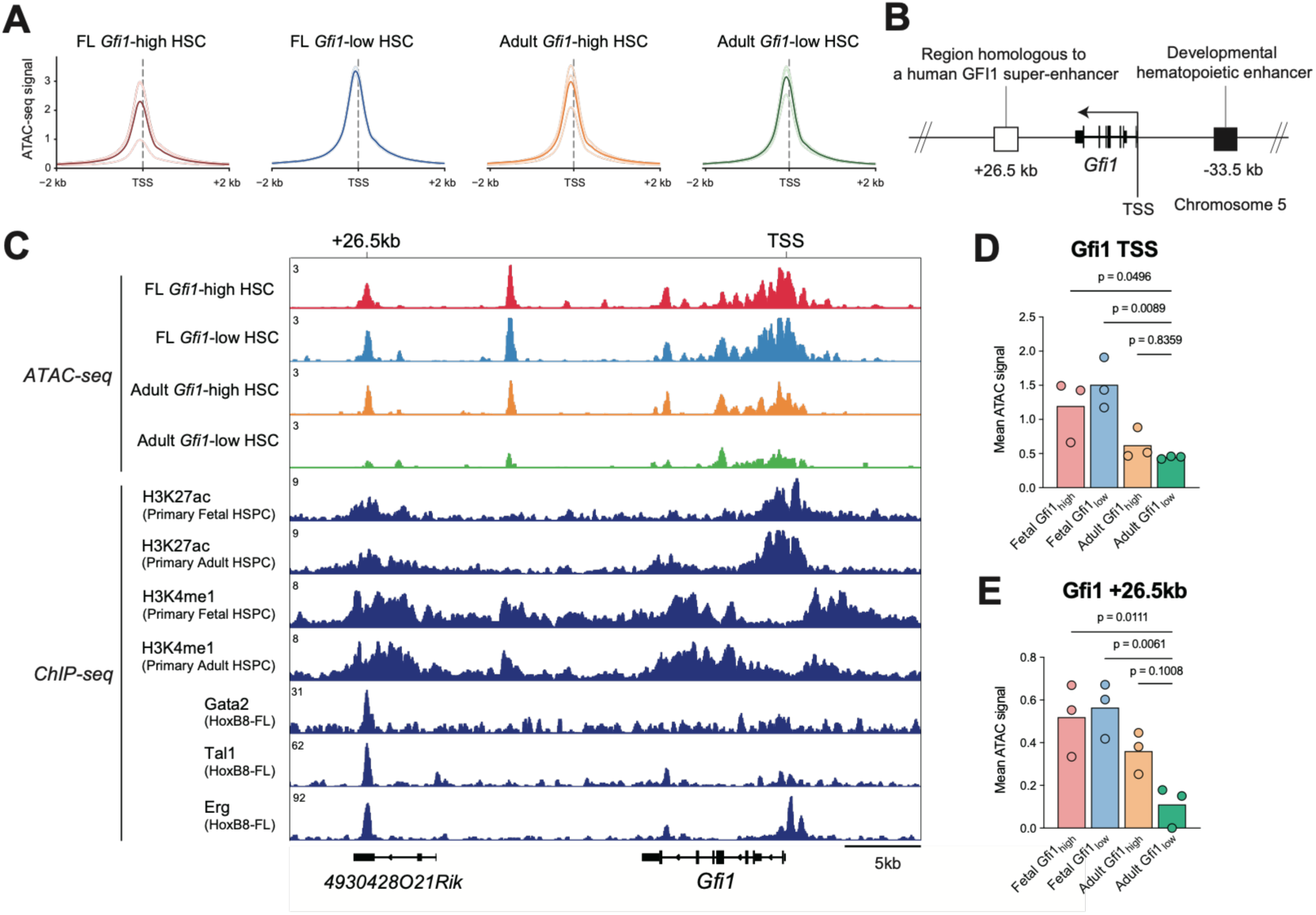
Cis-regulatory elements at the *Gfi1* locus are developmentally remodeled in HSCs. **(A)** ATAC-seq profiles centered on transcription start sites (TSSs; ±2 kb) in *Gfi1*-high and *Gfi1*-low HSCs isolated from E19.5 FL and adult BM. Thin lines represent individual biological replicates, and thick lines represent the mean profile for each group. **(B)** Schematic of the mouse *Gfi1* locus showing the -33.5 kb early hematopoietic enhancer, and the conserved +26.5 kb putative enhancer, which is orthologous to a functional enhancer within the human *GFI1* super-enhancer characterized in human erythroid leukemia cells. **(C)** Mean CPM-normalized ATAC-seq tracks spanning the *Gfi1* TSS and +26.5 kb putative enhancer in *Gfi1*-high and *Gfi1*-low HSCs isolated from E19.5 FL and adult BM, together with H3K27ac and H3K4me1 ChIP-seq profiles from fetal and adult HSPCs (GSE128760^14^) and GATA2, TAL1, and ERG ChIP-seq profiles from Hoxb8-FL cells (GSE84328^21^). **(D, E)** Quantification of mean CPM-normalized ATAC-seq signals at the *Gfi1* +26.5 kb putative enhancer **(D)** and TSS **(E)**.

## Discussion

In this study, we generated a physiological *Gfi1* knock-in reporter mouse to track the developmental dynamics of endogenous *Gfi1* expression. Using this model, we show that *Gfi1* reporter activity selectively marks ongoing EHT in vivo, reveals vascular-bed-specific differences in embryonic hemogenic activation, and identifies a functionally potent adult HSC subset that retains *Gfi1* expression.

A major strength of our *Gfi1*-T2A-mScarlet model is its high specificity for ongoing EHT. Although multiple transcription factors regulate EHT and HSC emergence ^23–34^, existing reporters are not strictly EHT-specific at E10.5. *Runx1, Gata2, and Evi1* reporters mark hemogenic endothelium and IAHCs, but also label non-EHT tissues within the embryonic trunk^23,24,27–29,35,36^. Previous *Gfi1* reporter studies linked *Gfi1* expression to EHT and showed that GFI1 and GFI1B promote hematopoietic emergence through LSD1/CoREST-mediated repression of the endothelial program^8,37^. This regulatory axis is also supported by zebrafish studies^38^. More recently, NOTCH1–JAG1 cis-inhibition was identified in *Gfi1*⁺ IAHCs as a mechanism that preserves HSC fate by limiting premature proliferation and differentiation^39^. Using our non-disruptive reporter, we found that nearly all c-Kit⁺ cells within IAHCs in the embryonic trunk were reporter-positive, whereas reporter activity was virtually absent from other major tissues. Thus, physiological *Gfi1* expression is tightly coupled to active EHT *in vivo*, supporting the use of this reporter to isolate EHT-competent cells and refine *ex vivo* strategies for generating functional HSCs^40,41^.

Previous whole-embryo three-dimensional analyses showed that c-Kit⁺ cells first appear in the VA and UA at around the 20-somite-pair (sp) stage^13^. These cells were absent from the DA until the 24/25-sp stage, suggesting that hemogenic activity begins earlier in the VA and UA^13,42^. Consistent with this, previous Runx1-based studies showed that Runx1 expression is prominent in the VA and UA at E9.5 and subsequently becomes detectable in the DA^24,43^. Because GFI1 acts downstream of RUNX1 during EHT^7,37^, earlier RUNX1 activation in the UA and VA may induce a GFI1-dependent hemogenic program in these arteries. However, previous studies primarily mapped c-Kit⁺ cells, RUNX1-positive regions, or accumulated clusters rather than directly visualizing the initiation of EHT itself. In this study, mScarlet-positive endothelial cells were markedly more abundant in the UA and VA than in the DA at E9.5, linking the earlier appearance of hematopoietic cells in these vessels to earlier and/or more frequent activation of the EHT-associated program.

Our study also highlights an important conceptual distinction between loss-of-function models and analysis of physiological expression states. Constitutive *Gfi1* knockout mice exhibit increased cycling and impaired quiescence in adult HSCs, accompanied by reduced long-term repopulating and self-renewal capacities^44,45^. However, because GFI1 is required from embryonic HSC emergence through adulthood, conventional germline knockout models cannot distinguish whether these phenotypes reflect defects in developmental HSC ontogeny, adult HSC maintenance, or both^7,8^. Moreover, complete gene deletion may not capture the biological significance of physiological differences in *Gfi1* expression. By contrast, our model preserves endogenous GFI1 function and enables functional analysis of HSC subsets defined by physiological Gfi1 expression levels. Recent dose-dependent analyses further support the idea that *Gfi1* expression levels influence HSC engraftment and differentiation capacity^46^.

The functional significance of *Gfi1* expression may be context-dependent across HSC ontogeny. FL HSCs occupy a distinct developmental state in which active proliferation coexists with robust self-renewal and repopulating capacity, whereas adult BM HSCs are reprogrammed toward long-term maintenance through deeper quiescence^1,17,47–55^. Fetal HSC identity has been linked to developmental regulators such as the *Lin28b*-*let-7*-*Hmga2* axis, *Sox17*-dependent programs, IGF2BP family members, and imprinted gene networks^56–61^. In our dataset, the *Gfi1*-high adult HSC fraction was enriched for multiple FL HSC-associated genes, including *H19*, *Igf2*, *Meg3*, *Mirg*, and *Igf2bp2*. Many of these genes, like *Gfi1* itself, are normally downregulated during fetal-to-adult transition. *Mirg* and *Meg3* reside within the Dlk1-Gtl2 locus, whereas *H19* and *Igf2* belong to the H19-Igf2 locus and both loci have been implicated in fetal HSC programs that support stem cell maintenance in a proliferative state ^62–64^. Together, these observations suggest that *Gfi1*-expressing adult BM HSCs partially retain a fetal-associated developmental program marked by imprinting-related genes. Rather than merely marking quiescence, residual *Gfi1* expression may characterize an adult LT-HSC state that reconciles a dormant transcriptional program with preserved self-renewal and regenerative capacity. Consistent with this idea, high *GFI1* expression marks stem-like memory CD8^+^ T cells and supports their long-term persistence by preserving proliferative potential^65^. However, our study does not determine whether Gfi1 directly regulates these imprinting-associated genes or whether this imprinting-related signature is mechanistically linked to the enhanced functional activity of *Gfi1*-high HSCs.

ATAC-seq analysis suggested that developmental *Gfi1* downregulation in HSCs may involve region-specific changes in chromatin accessibility at the *Gfi1* locus. Accessibility at the conserved +26.5-kb putative enhancer was reduced in adult *Gfi1*-low HSCs, whereas the −33.5-kb early hematopoietic enhancer^19^ showed no comparable reduction, indicating stage-specific regulation of distinct *cis*-regulatory elements. The +26.5-kb region is orthologous to a functional element within the human GFI1 super-enhancer and contains predicted RUNX1, GATA1/2, ERG, and TAL1 motifs^20^. Public ChIP-seq data^21^ from mouse Hoxb8-FL cells further showed occupancy of this region by GATA2, TAL1, ERG, RUNX1, FLI1, and other hematopoietic transcription factors. In human acute erythroid leukemia cells, the corresponding region was required for LSD1 inhibitor-induced GFI1 expression and reporter activity^20^. Together, its evolutionary conservation, motif composition, transcription factor occupancy, and functional validation support its designation as a conserved *Gfi1* cis-regulatory element whose reduced accessibility may contribute to fetal-to-adult *Gfi1* downregulation.

In summary, we establish a physiological *Gfi1* knock-in reporter that selectively marks ongoing EHT and reveals temporal differences in hemogenic activation across embryonic vascular beds. Developmental and functional analyses further identified an adult HSC subset characterized by relatively higher *Gfi1* expression, enhanced repopulating capacity, and fetal-associated transcriptional features. Together, these findings link developmental *Gfi1* dynamics to stage-specific HSC function and establish this reporter as a tool for isolating EHT-competent cells and investigating HSC heterogeneity.

## STAR Methods

### Mice

All mice used in this study were C57BL/6 genetic background. Mice were maintained under specific pathogen-free conditions. C57BL/6-Ly5.2 and C57BL/6-Ly5.1 mice were purchased from Japan SLC and Sankyo-Lab Service, respectively. Adult mice were analyzed at 8-12 weeks of age unless otherwise specified. Embryos were obtained by timed matings and staged according to embryonic day, somite pair number, and Theiler criteria in the eMouseAtlas resource (http://www.emouseatlas.org/emap/home.html). Gfi1-T2A-mScarlet mice were used as heterozygotes unless otherwise indicated. All animal experiments were approved by the Institutional Animal Care and Use Committee of Kumamoto University and conducted in accordance with institutional guidelines.

### Generation of Gfi1-T2A-mScarlet knock-in mice

*Gfi1*-T2A-mScarlet knock-in mice were generated by CRISPR/Cas9-mediated genome editing. A guide RNA targeting the sequence 5′-ATGGACTCAAATGAGTACCC-3′, followed by the PAM sequence 5′-TGG-3′, was used; this target sequence spans the 3′ end of the *Gfi1* coding sequence, the endogenous stop codon, and adjacent downstream sequence. The donor vector was designed in-house to insert a T2A-mScarlet cassette at the 3′ end of the *Gfi1* coding sequence, before the endogenous stop codon, by homology-directed repair. The cassette was flanked by 1,500-bp left and right homology arms corresponding to sequences surrounding the intended insertion site in the mouse *Gfi1* locus. The T2A sequence was included to allow Gfi1 and mScarlet to be translated as separate proteins from the endogenous *Gfi1* transcript. The designed donor sequence was synthesized and cloned into a pUC-based plasmid backbone by VectorBuilder (Yokohama, Japan). Pronuclear injection was performed as described previously^66^. Specifically, crRNA and tracrRNA (Sigma Aldrich) were diluted with nuclease-free water. The mixture was denatured at 95°C and then annealed by gradually cooling to room temperature (∼1 hour). The gRNA was mixed with Cas9 solution Cas9 protein (Thermo Fisher Scientific, A36497, 27 ng/µL final concentration) and TE buffer (10 mM Tris-HCl, 0.1 mM EDTA, pH 7.4). After incubation at 37℃ for 5 minutes, the mixture was mixed with guide RNA components consisting of crRNA and tracrRNA, and the donor plasmid (10 ng/µL final concentration) and were microinjected into fertilized one-cell embryos from C57BL/6J mice. Cas9 protein, guide RNA components consisting of crRNA and tracrRNA, and the donor plasmid were microinjected into fertilized one-cell embryos from C57BL/6J mice. Injected embryos were transferred into pseudo-pregnant females together with ICR embryos to support implantation. Founder candidates were selected from black pups derived from the C57BL/6J background. Founder candidates were screened by PCR to confirm 5′- and 3′-junction integration, distinguish knock-in and wild-type alleles, and exclude donor plasmid backbone integration. Correctly targeted founders showed amplification of both 5′ and 3′ junctions and no detectable donor backbone signal. Primer sequences are listed in Supplementary Table 1. Confirmed founder mice were crossed with wild-type C57BL/6J mice to establish the *Gfi1*-T2A-mScarlet knock-in mice. For genotyping of F2 and later generations, a three-primer PCR was used to distinguish wild-type, heterozygous knock-in, and homozygous knock-in mice. The knock-in and wild-type alleles produced 564-bp and 330-bp amplicons, respectively. Primer sequences are listed in Supplementary Table 1. All experiments were performed using F2 or later generations.

### Whole-mount immunostaining of embryos

Whole-mount immunostaining was performed based on a previously described protocol ^67^. Briefly, embryos were fixed in 2% paraformaldehyde/PBS on ice for 25 min, followed by dehydration in 50% methanol/PBS for 10 min and 100% methanol three times for 10 min each. Embryos were rehydrated into PBS, blocked in PBS-MT (PBS containing 0.4% Triton X-100 and 1% skim milk) for 1 hour, and incubated overnight at 4°C with primary antibodies diluted in PBS-MT. After primary antibody incubation, samples were washed three times with PBS-MT for 1 hour each. Secondary antibodies diluted in PBS-MT were then applied overnight at 4°C. Samples were washed once with PBS-MT and twice with PBS-T (PBS containing 0.4% Triton X-100) for 1 hour each, followed by dehydration in 50% methanol/PBS and 100% methanol for approximately 5 min each. The primary antibodies used were to c-Kit (rat, clone 2B8; eBioscience), CD31 (hamster clone 2H8; Invitrogen), CD31 (rat clone MEC13.3; BD Pharmingen), RFP (Rockland Immunochemicals, Cat. No. 600-401-379), Gfi1 (rabbit clone E5J6J; Cell Signaling Technology), and IgG (rabbit clone DA1E; Cell Signaling Technology). The secondary antibodies used were goat anti-hamster IgG-Alexa Fluor 488 (Jackson ImmunoResearch), donkey anti-rat IgG-Cy3 (Jackson ImmunoResearch), donkey anti-rabbit IgG-Cy3 (Jackson ImmunoResearch), donkey anti-rat IgG-Alexa Fluor 647 (Invitrogen), goat anti-rabbit IgG-Alexa Fluor 647 (Invitrogen).

### Confocal microscopy and image analysis

After immunostaining, embryo proper and yolk sac samples were optically cleared in a 1:2 benzyl alcohol/benzyl benzoate mixture and imaged by confocal microscopy. Images were acquired on an Olympus FV1200 confocal microscope equipped with GaAsP PMT detectors, a UPLSAPO 20×/NA 0.75 objective, and FV10-ASW software. Large embryonic regions were imaged by tile scanning with a motorized x-y stage. Three-dimensional images were reconstructed from z-stack datasets using Olympus FLUOVIEW software.

### Preparation of single-cell suspensions from embryos

Embryonic bodies were collected in differentiation medium consisting of αMEM supplemented with 10% FCS and 1% penicillin/streptomycin. Tissues were enzymatically digested with 1 mg/ml Dispase and Collagenase at 37°C for 30 min, with gentle tapping every 10 min. Digestion was stopped by adding excess differentiation medium, and tissues were mechanically dissociated by gentle pipetting. Cell suspensions were passed through a nylon mesh to remove aggregates, washed, and incubated with differentiation medium at 37°C for 30 min to allow antigen recovery.

### Flow cytometric analysis and sorting of bone marrow and peripheral blood cells

Bone marrow cells were obtained by gently crushing the pelvis, femurs, and tibias in Dulbecco’s Modified Eagle Medium (Sigma) supplemented with 10% fetal bovine serum (Biowest). After red blood cell lysis with ACK buffer (Thermo Fisher Scientific), the cell suspensions were passed through 100-μm strainers and washed once with Dulbecco’s phosphate-buffered saline (PBS; Sigma) containing 2% FBS. For HSC enrichment, BM cells were magnetically labeled with microbead-conjugated anti-c-Kit antibodies (Miltenyi Biotec) and separated using the autoMACS Pro Separator (Miltenyi Biotec). Peripheral blood cells were treated with ACK buffer before antibody staining. Cells were stained with fluorescence-conjugated antibodies: c-Kit (clone 2B8), CD150 (TC15-12F12.2), CD48 (HM48-1), EPCR (eBio1560, eBioscience), CD41 (MWReg30), Sca-1 (D7), Ter-119 (TER-119), CD4 (GK1.5), CD8a (53-6.7), B220 (RA3-6B2), Gr-1 (RB6-8C5), Mac-1/CD11b (M1/70), IgD (11-26, eBioscience), IgM (II/41, eBioscience), CD3e (145-2C11), CD45 (30-F11, BD Biosciences), Ly6C (HK1.4), Ly6G (1A8), and VE-cadherin/VECAD (BV13). All antibodies were purchased from BioLegend unless otherwise indicated. Cells were stained in FACS buffer containing 2% FBS in PBS, washed, and resuspended with propidium iodide for dead-cell exclusion. Flow cytometric analysis and sorting were performed using a FACSAria III cell sorter and a BD FACSymphony cell analyzer (BD Biosciences), and data were analyzed with FlowJo v10.10.1 (Beckman Coulter). Cell populations were defined as follows: KSL cells, Lineage⁻c-Kit⁺Sca-1⁺; MPP2, CD150⁺CD48⁺ KSL; MPP3/4, CD150⁻CD48⁺ KSL; ST-HSCs, CD150⁻CD48⁻ KSL; LT-HSCs/HSCs, CD150⁺CD48⁻ KSL; and LK cells, Lineage⁻c-Kit⁺Sca-1⁻. For adult samples, lineage-positive cells were defined using antibodies against CD4, CD8a, Mac-1/CD11b, Gr-1, B220, and Ter-119. For fetal samples, lineage-positive cells were defined using antibodies against CD4, CD8a, Gr-1, B220, and Ter-119, excluding Mac-1/CD11b from the lineage cocktail because fetal hematopoietic stem and progenitor cells can express Mac-1/CD11b^55,68^.

### Competitive transplantation assays

For transplantation assays, CD150⁺CD48⁻KSL LT-HSCs were FACS-purified from E14.5 fetal liver or adult bone marrow of *Gfi1*-T2A-mScarlet reporter mice and divided into *Gfi1*-high and *Gfi1*-low fractions, defined as the top 30% and bottom 30% of cells based on mScarlet fluorescence intensity, respectively. Sorted *Gfi1*-high or *Gfi1*-low CD45.2⁺ LT-HSCs were co-transplanted with 2 × 10⁵ CD45.1⁺ competitor BM cells into lethally irradiated CD45.1⁺ recipients. Donor-derived reconstitution was assessed by peripheral blood CD45.2 chimerism at 4-week intervals up to 16 weeks after transplantation.

For lineage analysis, peripheral blood cells were stained with antibodies against CD45.1, CD45.2, Mac-1, Gr-1, B220, CD4, and CD8. Myeloid cells were defined as Mac-1⁺ and/or Gr-1⁺ cells, B cells as B220⁺ cells, and T cells as CD4⁺ or CD8⁺ cells. Antibodies used included CD45.1 (A20, FITC), CD45.2 (104, APC), Mac-1 (M1/70, PerCP-Cy5.5), Gr-1 (RB6-8C5, PE), B220 (RA3-6B2, PE-Cy7), CD4 (GK1.5, APC-Cy7), and CD8a (53-6.7, APC-Cy7), all purchased from BioLegend. Cells were analyzed using a FACSAria III. For secondary transplantation, 5 × 10⁶ whole BM cells harvested from primary recipient mice 16 weeks after transplantation were transplanted into lethally irradiated CD45.1⁺ secondary recipient mice. Donor-derived reconstitution was assessed based on CD45.2 expression.

### RNA sequencing

RamDA-seq libraries were prepared as previously described^69^. Briefly, 100 *Gfi1*-high or *Gfi1*-low FACS-purified HSCs were sorted directly into lysis buffer, treated with DNase I, and subjected to first- and second-strand cDNA synthesis using NSR/oligo-dT primers and Klenow fragment. Purified double-stranded cDNA was converted into sequencing libraries using Nextera XT-based tagmentation and indexing. Libraries were sequenced on an NovaSeq X Plus to generate paired-end reads. Sequencing reads were mapped to the mouse reference genome (mm10) using HISAT2 ^70^. Gene raw counts were then generated using featureCounts. The resulting count matrices were imported into RNAseqChef ^71^, and differential gene expression analysis was performed using DESeq2. Pre-ranked GSEA was then performed using clusterProfiler, with genes ranked by log2 fold change. The ranked list was oriented so that positive values corresponded to Gfi1-high HSCs. Custom HSC signatures, including LT-HSC ^15^, dormant HSC, primed HSC ^16^, and fetal liver HSC signatures ^17^, were tested, and enrichment was evaluated by normalized enrichment score and P value.

### Single-cell RNA-seq reanalysis

Public single-cell RNA-seq data^14^ (GSE128761) were analyzed using Seuratv5.4.0. Gene-barcode matrices were imported using the “Read10X” function, and Seurat objects were generated using the “CreateSeuratObject” function. Quality filtering for each cell was performed based on the following criteria: nFeature_RNA > 200, nFeature_RNA < 6,000, and percent mitochondrial genes < 10%. Feature counts were log-normalized using the “NormalizeData” function with a scale factor of 10,000. Two thousand highly variable features were selected for PCA. Clustering was performed using “FindNeighbors” and “FindClusters” with the first 10 principal components at a resolution of 0.5. UMAP visualization was performed using the same principal components. Gene expression patterns were visualized using “FeaturePlot” and “VlnPlot”. For adult HSC subset analysis, adult HSCs were extracted and reanalyzed separately. They were reclustered at a resolution of 0.3 and visualized by UMAP using the first 20 principal components. A dormant HSC-derived module score was calculated using “AddModuleScore”, and cells were stratified into three groups according to their HSC score ^16^. Gene expression patterns were compared among the HSC-score groups.

### ATAC-seq library preparation, sequencing, and data processing

For each sample, 1,000 FACS-purified HSCs were washed with ice-cold PBS and lysed in ATAC resuspension buffer containing 0.1% IGEPAL CA-630, 0.1% Tween-20, and 0.01% digitonin for 5 min on ice. Nuclei were washed and subjected to transposition with TD buffer and TDE1 Tn5 transposase (Illumina) in the presence of 0.1% Tween-20 and 0.01% digitonin at 37°C for 30 min with agitation at 900 rpm. DNA was purified using a MinElute Reaction Cleanup Kit (QIAGEN). Libraries were generated by PCR amplification, with the optimal cycle number determined by quantitative PCR. Library fragment size and concentration were assessed using a 4150 TapeStation System (Agilent Technologies). Pooled libraries were sequenced on a NextSeq 500 system (Illumina) to generate 75-bp single-end reads. Raw FASTQ files were assessed using FastQC, and adapter sequences were removed using “Trim Galore!”. Trimmed reads were aligned to the mm10 mouse reference genome using Bowtie2. CPM-normalized genome-wide coverage tracks were generated in BigWig format using bamCoverage.

### Statistical analysis

Statistical analyses were performed using Prism version 11 software (GraphPad). Differences between two groups were assessed using Welch’s t test. using one-way ANOVA followed by Dunnett’s multiple-comparisons test or Brown-Forsythe and Welch ANOVA. Data are presented as mean ± SEM unless otherwise indicated. Two-tailed P values are shown, with statistical significance indicated as follows: *p < 0.05; **p < 0.01; ***p < 0.001; ****p < 0.0001; n.s., not significant.

## Acknowledgments

We thank Miho Kataoka and Kenta Kikuchi for their excellent technical assistance. We appreciate the technical support provided by the core facilities of the International Research Center for Medical Sciences (IRCMS), Kumamoto University, and the excellent animal care provided by the Center for Animal Resources and Development (CARD), Kumamoto University. We also thank the FACS Core and the Mouse Core at the Institute of Medical Science, the University of Tokyo.

This research was supported by a JSPS Grant-in-Aid for Scientific Research (S) (TS, 18H05284), the Chinese Academy of Medical Sciences Innovation Fund for Medical Sciences (TS, 2024-12M-3-017), and the National Natural Science Foundation of China (TS, W2441024). This work was also supported by JSPS Grants-in-Aid for Young Scientists (TY, 24K19227 and 26K19430), a JSPS Grant-in-Aid for Challenging Research (Exploratory) (YT, 24K11520), a Medical Research Grant from the Takeda Science Foundation (YT, 2023044429), a research grant from the Mochida Memorial Foundation for Medical and Pharmaceutical Research (TY and YT), a Research Encouragement Grant from the Novartis Foundation (Japan) for the Promotion of Science (YT), the General Research Award (Credit Saison Award) from the Japan Leukemia Research Fund (YT), a research grant from the Shinnihon Foundation of Advanced Medical Treatment Research (TY and YT), a grant from the SENSHIN Medical Research Foundation (TY and YT), a LEGEND Research Grant from BioLegend (TY), a Research Grant from the Japanese Society of Hematology (TY and YT), and the Kanehara Ichiro Award (TS). This research was also supported by the MEXT Promotion of Distinctive Joint Usage/Research Center Support Program (Grant Number JPMXP0724020288) at the Advanced Medical Research Center, Yokohama City University.

## Authorship Contributions

T.Yabushita. and Y.T. conceived the project, designed and performed most of the experiments, analyzed and interpreted the data, and wrote the manuscript. T.F., T.I., T.I., K.W., K.K., M.T., S.K.M., T.Yokomizo., T.N., T.U, A.N., D.K., S.Y., H.T., T.T, and M.O. assisted with the experiments.

T.Yokomizo., T.N., T.U, A.N., D.K., S.Y., H.T., T.T, and M.O. advised on the data interpretation, discussed and made suggestions for this study. T.S. supervised the project, interpreted the data, and participated in writing the manuscript.

