## Supplemental figures for "Developmental *Gfi1* Dynamics Define Hematopoietic Emergence and Adult Hematopoietic Stem Cell Potency"

#### Figure S1.

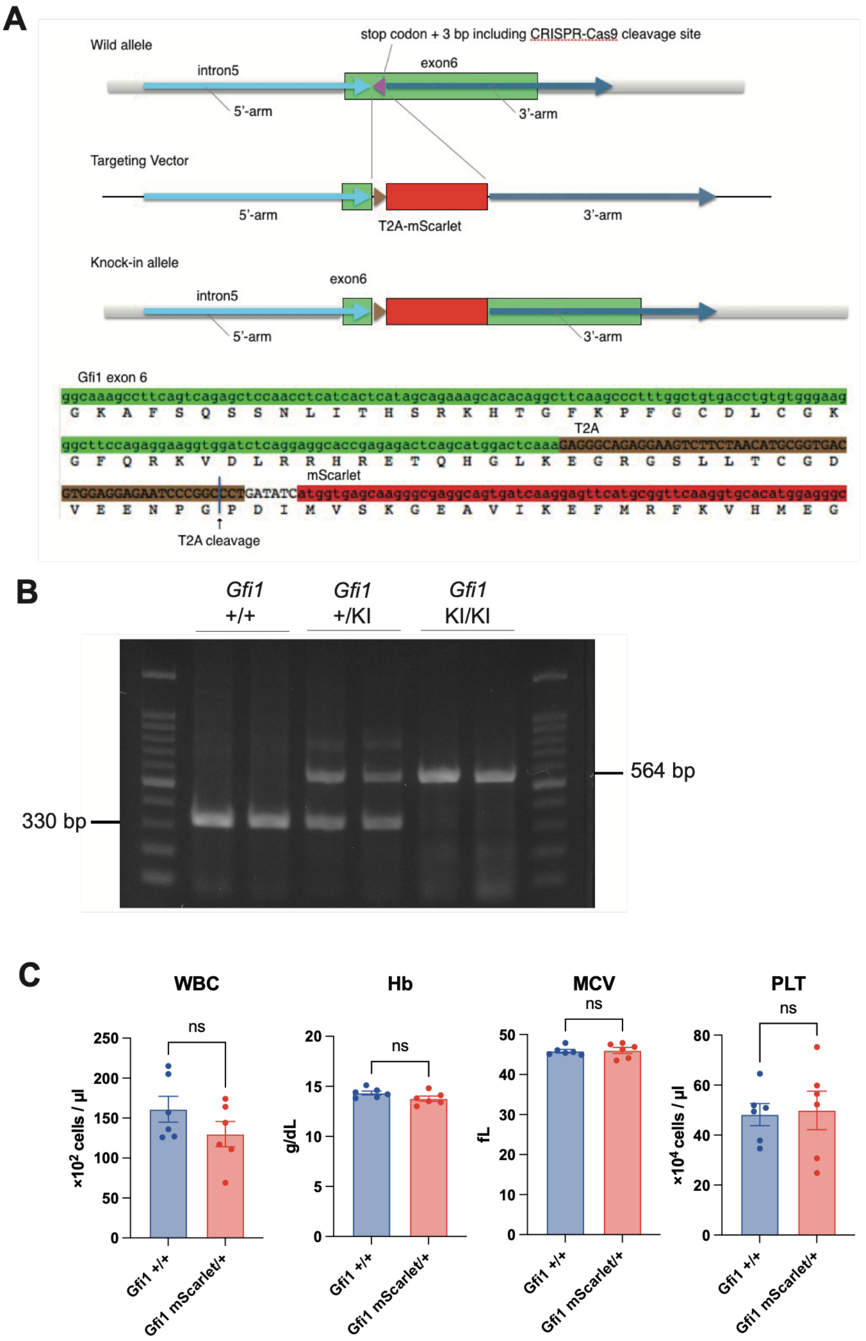


**Figure S1. Validation of the *Gfi1*-T2A-mScarlet knock-in allele**

**(A)** Schematic representations of the *Gfi1* wild-type allele, the CRISPR/HDR targeting vector, and the knock-in (KI) allele. A T2A–mScarlet cassette was inserted immediately upstream of the endogenous *Gfi1* stop codon using homology arms that span the Cas9 cleavage site. This gene editing preserves endogenous *Gfi1* expression, while mScarlet is co-translated as a faithful reporter of endogenous Gfi1 expression. **(B)** Genotyping PCR capable of simultaneously detecting wild-type and knock-in alleles. The wild-type allele yields a 330-bp PCR product, and the KI allele yields a 564-bp product, enabling discrimination of +/+ (WT/WT), mScarlet/+ (KI/WT), and mScarlet/mScarlet (KI/KI) genotypes. **(C)** Peripheral blood counts in WT and heterozygous KI mice.

#### Figure S2.

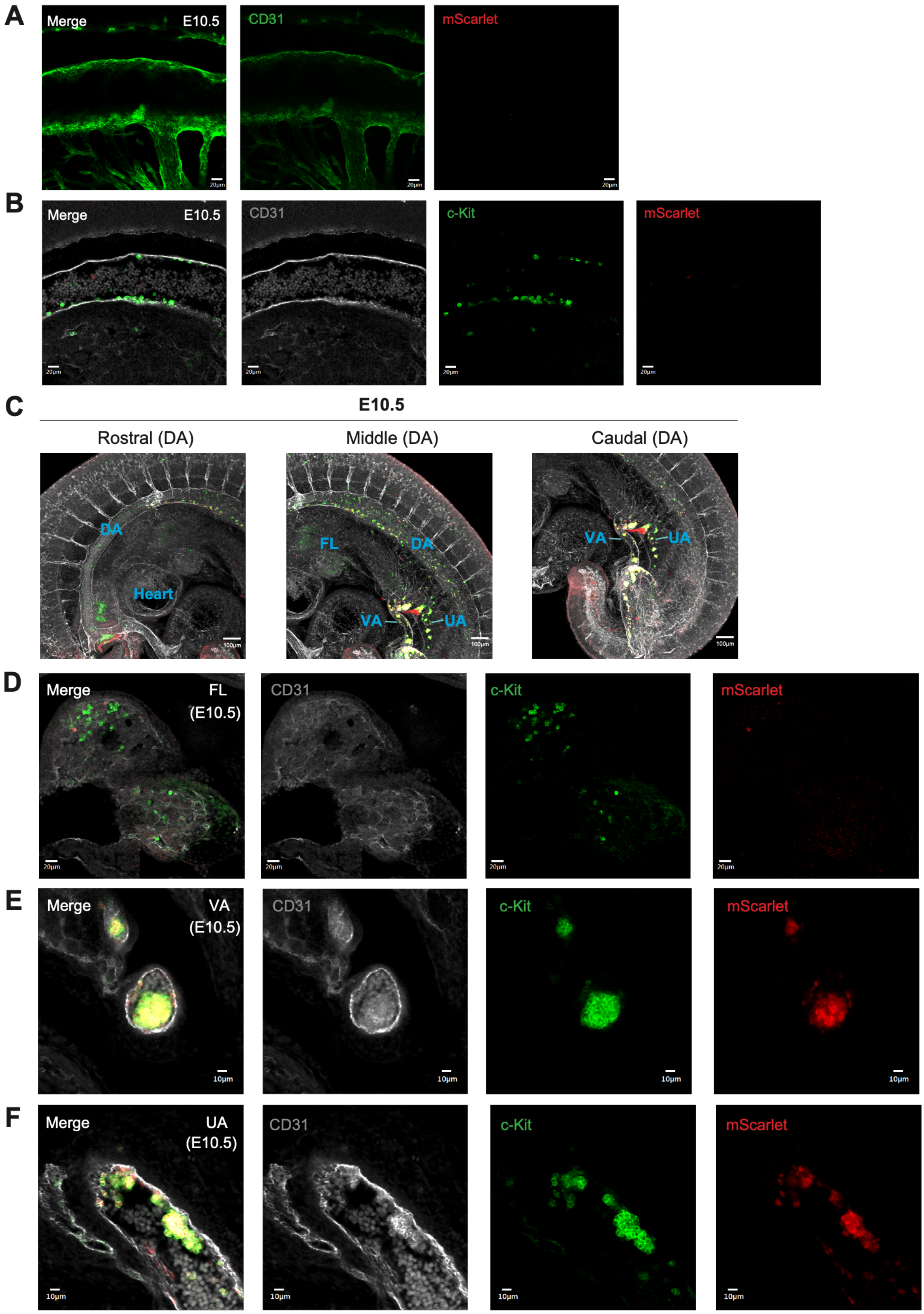
A
Merge
E10.5
CD31
mScarlet
B
Merge
E10.5
CD31
c-Kit
mScarlet
C
E10.5
Rostral (DA)
Middle (DA)
Caudal (DA)
DA
Heart
FL
DA
VA
UA
VA
UA
D
Merge
FL
(E10.5)
CD31
c-Kit
mScarlet
E
Merge
VA
(E10.5)
CD31
c-Kit
mScarlet
F
Merge
UA
(E10.5)
CD31
c-Kit
mScarlet

**Figure S2. *Gfi1*-mScarlet reporter activity is restricted to intra-aortic hematopoietic clusters and hematopoietic clusters in the umbilical and vitelline arteries.**

**(A, B)** Negative control images from E10.5 wild-type (WT) embryos. **(A)** Single-plane confocal images of the dorsal aorta (DA) and vitelline artery (VA), shown as merged, CD31, and mScarlet channels. **(B)** Single-plane confocal images of intra-aortic hematopoietic clusters (IAHCs) in the DA, shown as merged, CD31, c-Kit, and mScarlet channels. **(C)** Three-dimensional rendered confocal images of the rostral, middle, and caudal regions of the DA from E10.5 Gfi1-mScarlet KI/WT embryos. CD31 is shown in gray, c-Kit in green, and mScarlet in red. **(D–F)** Single-plane confocal images of the E10.5 Gfi1-mScarlet KI/WT fetal liver (FL) **(D)**, vitelline artery (VA) **(E)**, and umbilical artery (UA) **(F)**, shown as merged, CD31, c-Kit, and mScarlet channels. CD31 is shown in gray, c-Kit in green, and mScarlet in red.

#### Figure S3.

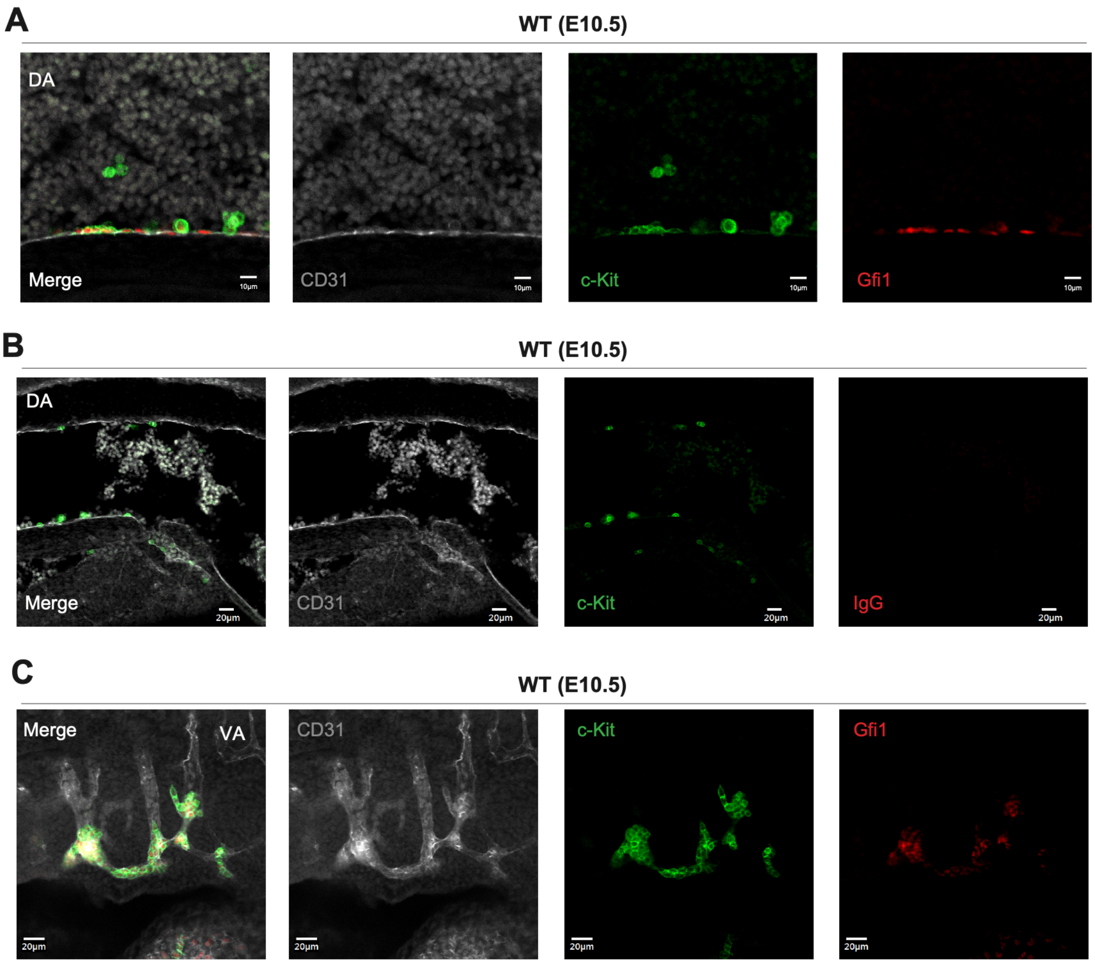


**Figure S3. Endogenous Gfi1 immunostaining confirms the distribution of *Gfi1*-mScarlet reporter activity in intra-aortic hematopoietic clusters and the umbilical/vitelline arteries.** All panels show single-plane confocal images. For each panel, images are presented in the following order: merged image (far left), CD31 (gray), c-Kit (green), and either Gfi1 or isotype control IgG (red), as indicated. **(A)** E10.5 wild-type (WT) ventral dorsal aorta (DA) immunostained for endogenous Gfi1. **(B)** E10.5 WT DA stained with isotype control IgG. **(C)** E10.5 WT vitelline artery (VA) immunostained for endogenous Gfi1.

#### Figure S4.

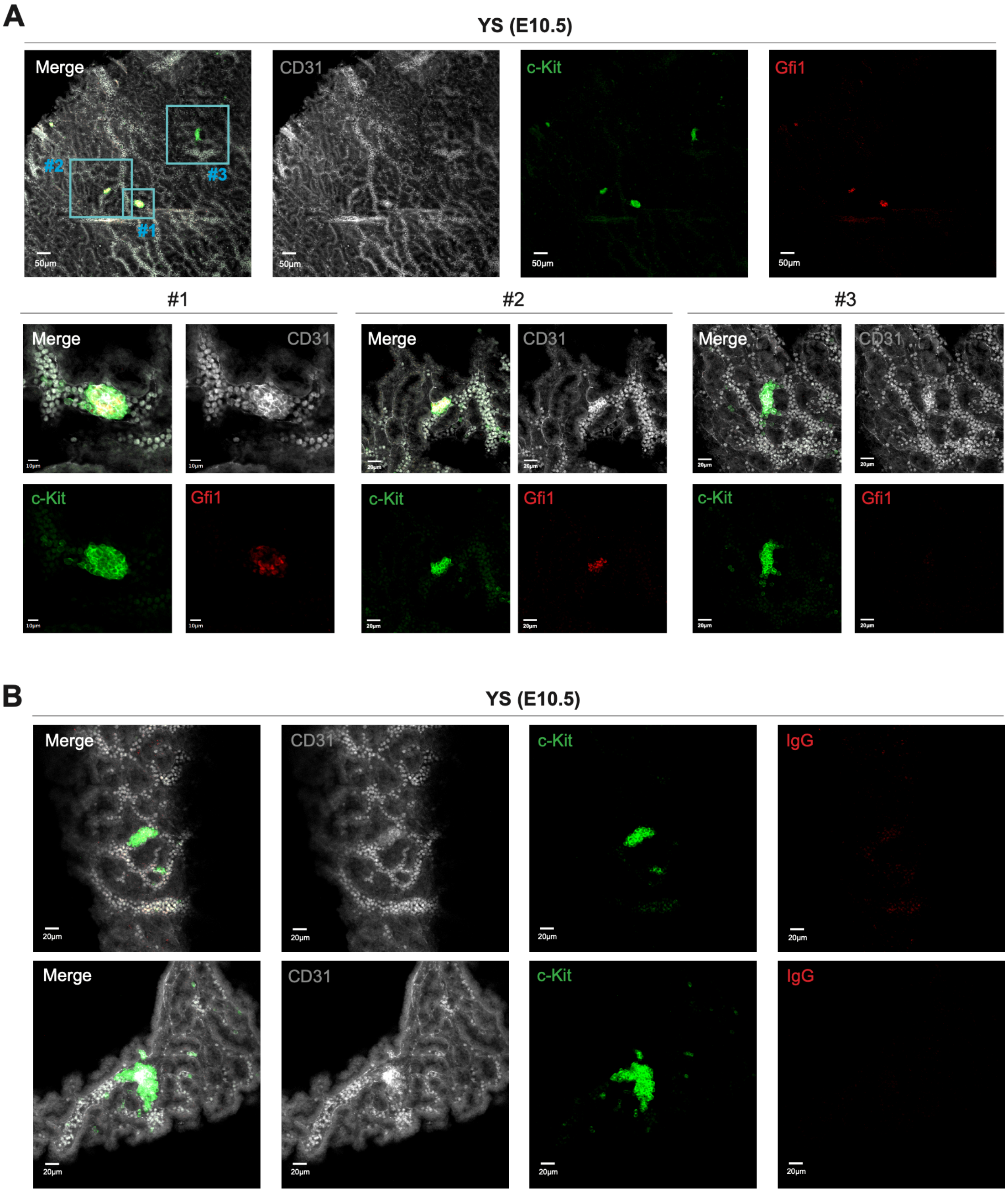


**Figure S4. Heterogeneous endogenous Gfi1 immunostaining in E10.5 yolk sac hematopoietic clusters.**

**(A)** Single-plane confocal images of the E10.5 wild-type (WT) yolk sac immunostained for endogenous Gfi1. Low-magnification views (upper) and high-magnification views (lower) of three c-Kit+ clusters (#1–#3) are shown, including Gfi1-positive clusters (#1 and #2) and a Gfi1-negative cluster (#3). CD31 is shown in gray, c-Kit in green, and Gfi1 in red. **(B)** Single-plane confocal images of the E10.5 WT yolk sac stained with isotype control IgG. CD31 is shown in gray, c-Kit in green, and IgG in red.

### Figure S5.

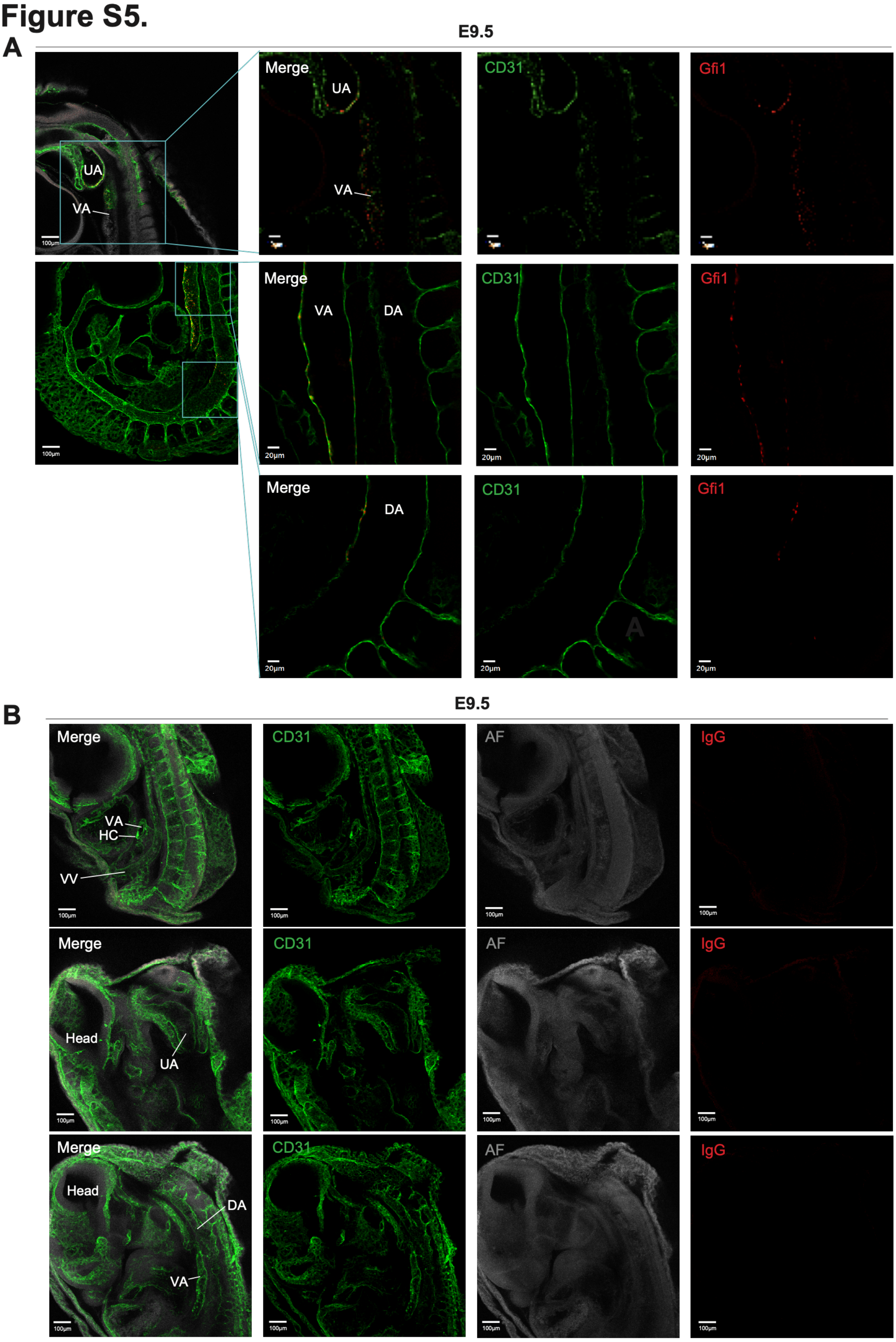
A
E9.5
UA
VA
Merge
UA
VA
CD31
Gfi1
Merge
VA
DA
CD31
Gfi1
Merge
DA
CD31
Gfi1
B
E9.5
Merge
VA
HC
VV
CD31
AF
IgG
Merge
Head
UA
CD31
AF
IgG
Merge
Head
DA
VA
CD31
AF
IgG

**Figure S5. Gfi1 immunostaining is preferentially detected in the embryonic umbilical and vitelline arteries at E9.5.**

**(A-B)** Single-plane confocal images of E9.5 wild-type embryos stained with CD31 (green) together with either anti-Gfi1 antibody (A, red) or isotype control IgG (B, red). UA, umbilical artery; VA, vitelline artery; VV, vitelline vein; DA, dorsal aorta; AF, autofluorescence.

#### Figure S6.

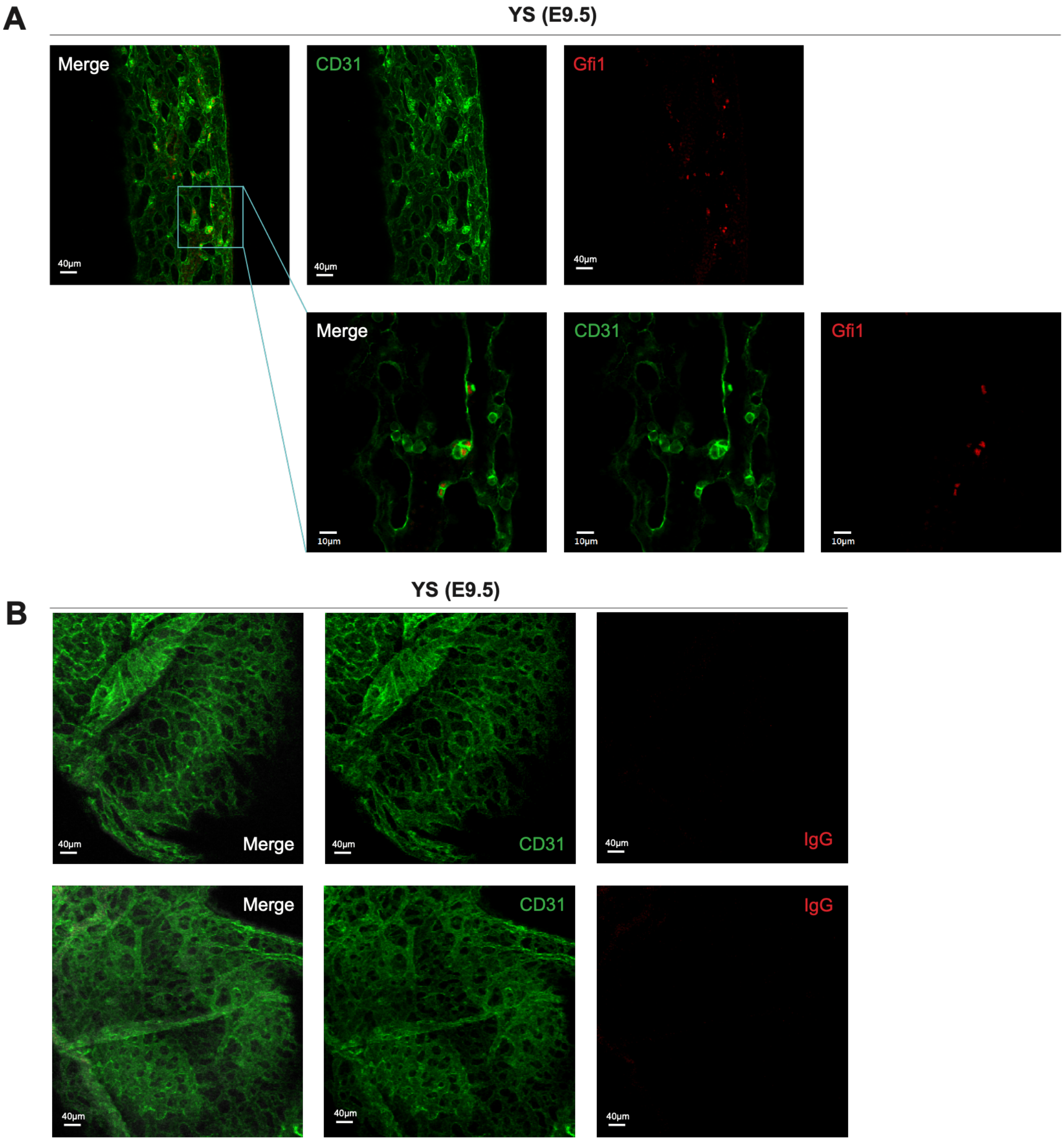


**Figure S6. Gfi1 labels hematopoietic clusters in the E9.5 yolk sac.**
**(A and B)** Single-plane confocal images of E9.5 wild-type yolk sac stained with CD31 (green) together with either Gfi1 antibody (A, red) or isotype control IgG (B, red).

**Figure S7.**

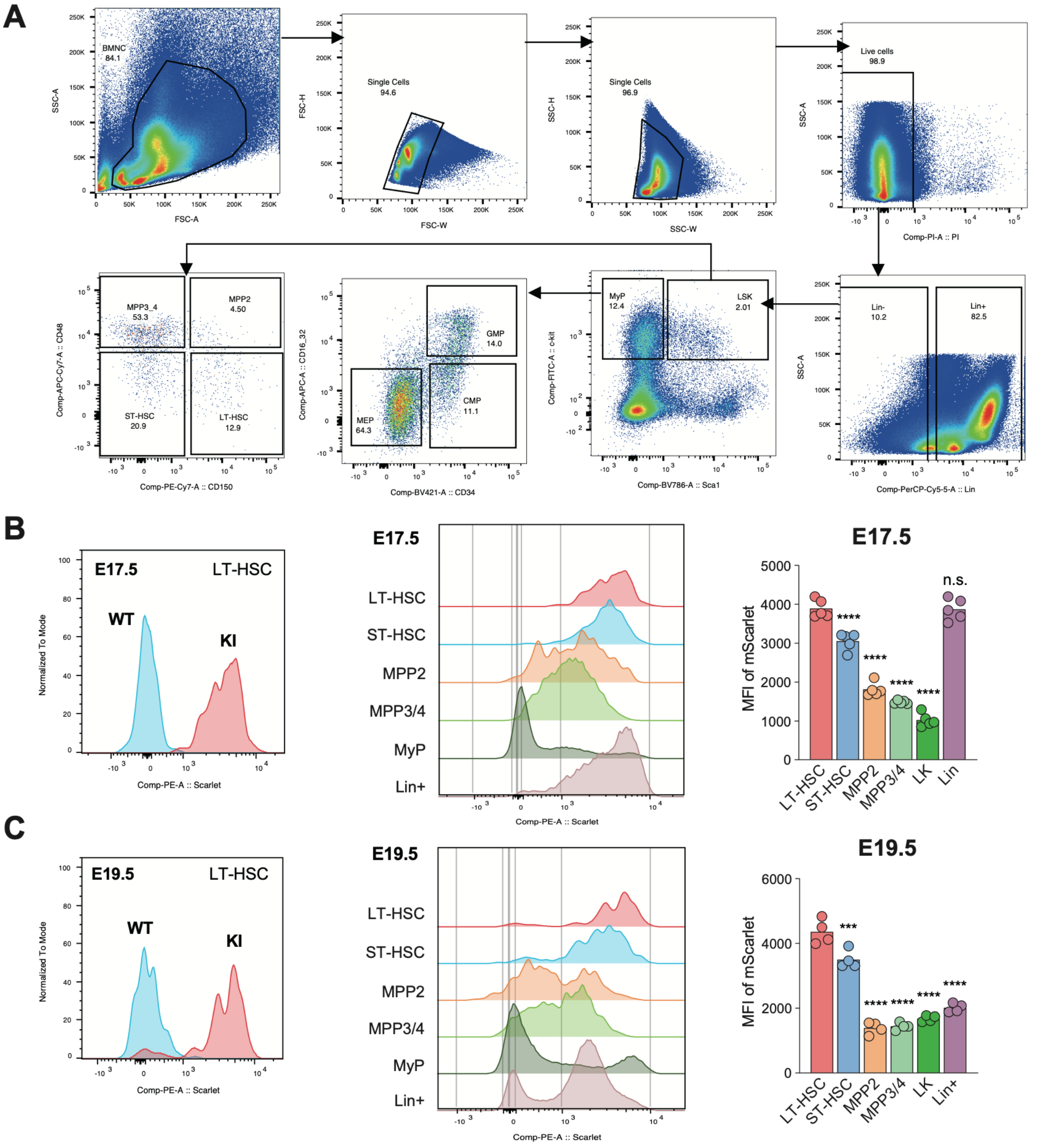


**Figure S7. Dynamic regulation of *Gfi1* reporter expression during fetal-to-adult hematopoietic maturation.**

**(A)** Flow cytometric gating strategy for myeloid progenitors (MyP; Lin⁻Sca-1⁻c-Kit⁺; LK fraction) and the LSK fraction (Lin⁻Sca-1⁺c-Kit⁺). MyP includes megakaryocyte–erythroid progenitors (MEP), common myeloid progenitors (CMP), and granulocyte–macrophage progenitors (GMP). LSK contains multipotent progenitors (MPP; comprising MPP2, MPP3, and MPP4 subsets), short-term HSCs (ST-HSCs), and long-term HSCs (LT-HSCs). **(B-C)** Representative histograms of mScarlet fluorescence in E17.5 (B) and E19.5 (C) fetal liver LT-HSCs (left; *Gfi1*–mScarlet⁺/WT in red; WT in blue). Middle panels show histograms across hematopoietic subsets (LT-HSC, ST-HSC, MPP2, MPP3/4, MyP, and Lin⁺), and right panels show mean fluorescence intensity (MFI) quantification. Both developmental stages exhibit a consistent hierarchy of reporter expression.

### Figure S8.

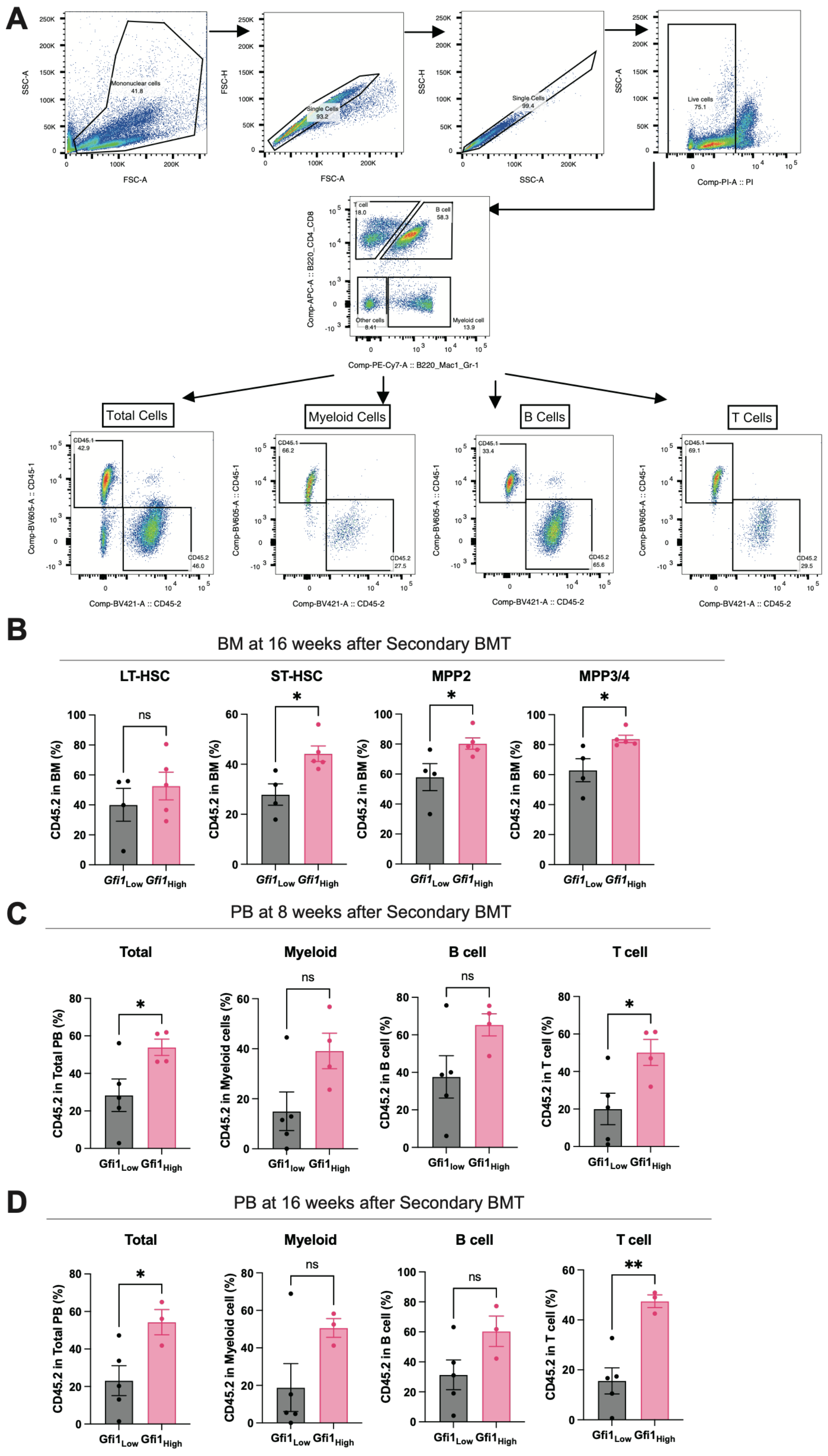

A
Total Cells
Myeloid Cells
B Cells
T Cells
B
BM at 16 weeks after Secondary BMT
LT-HSC
ST-HSC
MPP2
MPP3/4
C
PB at 8 weeks after Secondary BMT
Total
Myeloid
B cell
T cell
D
PB at 16 weeks after Secondary BMT
Total
Myeloid
B cell
T cell

**Figure S8. BM transplantation analysis of FL E14.5 HSCs.**
**(A)** Schematic representation of the strategy for peripheral blood chimerism analysis after transplantation. T cells were identified as $CD4^+$ or $CD8^+$ cells, B cells as $B220^+$ cells, and myeloid cells as $CD11b^+$ or $Gr\text{-}1^+$ cells. The frequencies of $CD45.2^+$ donor-derived cells and $CD45.1^+$ recipient-derived cells were assessed. **(B)** Bone marrow chimerism at 16 weeks primary transplantation of FL-HSCs. Donor chimerism was quantified in LT-HSC, ST-HSC, MPP2, and MPP3/4 fractions. **(C-D)** Peripheral blood chimerism at 8 weeks (C) and 16 weeks (D) after secondary transplantation of FL E14.5 HSCs shown for total cells, myeloid cells, B cells, and T cells.

### Figure S9.

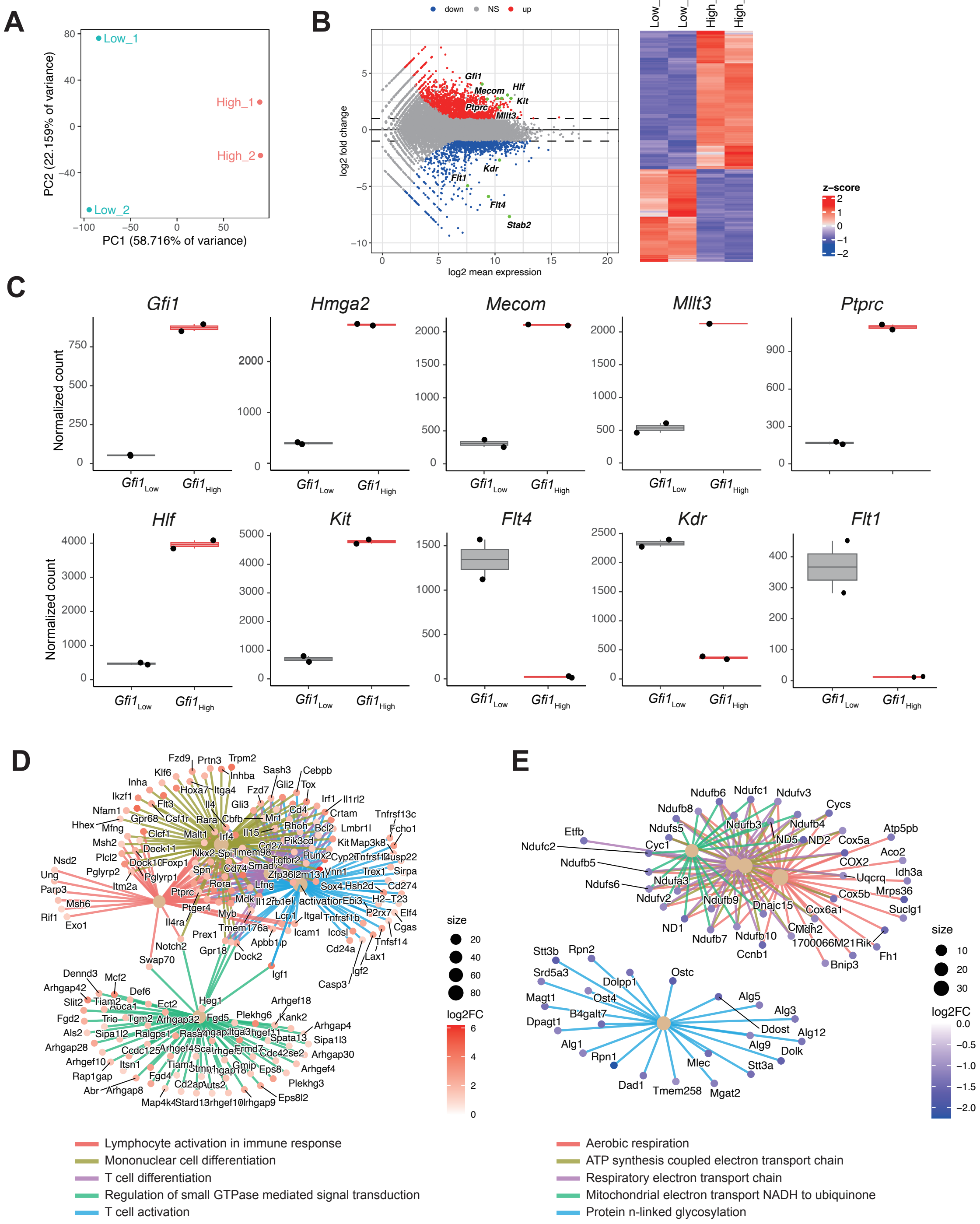
A
Low_1
High_1
High_2
Low_2
PC2 (22.159% of variance)
PC1 (58.716% of variance)
B
down
NS
up
log2 fold change
log2 mean expression
Gfi1
Mecom
Hlf
Kit
Ptprc
Mllt3
Kdr
Flt1
Flt4
Stab2
Low_1
Low_2
High_1
High_2
z-score
C
Gfi1
Hmga2
Mecom
Mllt3
Ptprc
Hlf
Kit
Flt4
Kdr
Flt1
Normalized count
Gfi1 Low
Gfi1 High
D
size
log2FC
Lymphocyte activation in immune response
Mononuclear cell differentiation
T cell differentiation
Regulation of small GTPase mediated signal transduction
T cell activation
E
size
log2FC
Aerobic respiration
ATP synthesis coupled electron transport chain
Respiratory electron transport chain
Mitochondrial electron transport NADH to ubiquinone
Protein n-linked glycosylation

**Figure S9. Transcriptomic distinctions between *Gfi1*-High and *Gfi1*-Low fetal liver HSCs in *Gfi1*-mScarlet E14.5 embryo.**

**(A)** Principal component analysis (PCA) plot of bulk RNA-seq profiles from *Gfi1*-high and *Gfi1*-low fetal liver (FL) HSCs at E14.5 (n = 2). **(B)** MA plot and hierarchical clustering heatmap showing differentially expressed genes (DEGs) between *Gfi1*-high and *Gfi1*-low FL HSCs (n = 2). DEGs were defined as genes with an adjusted *P* value $< 0.05$ and an absolute $\log_2$ fold change $> 2$. **(C)** Normalized RNA-seq read counts for representative genes (*Gfi1, Hmga2, Mecom, Mllt3, Ptprc, Hlf, Kit, Flt4, Kdr,* and *Flt1*) in *Gfi1*-high and *Gfi1*-low FL HSCs (n = 2). **(D, E)** Gene-concept network analyses showing selected enriched biological processes and contributing genes identified from the 1,684 genes preferentially expressed in *Gfi1*-high FL HSCs (D) and the 1,122 genes preferentially expressed in *Gfi1*-low FL HSCs (E).

**Figure S10.**

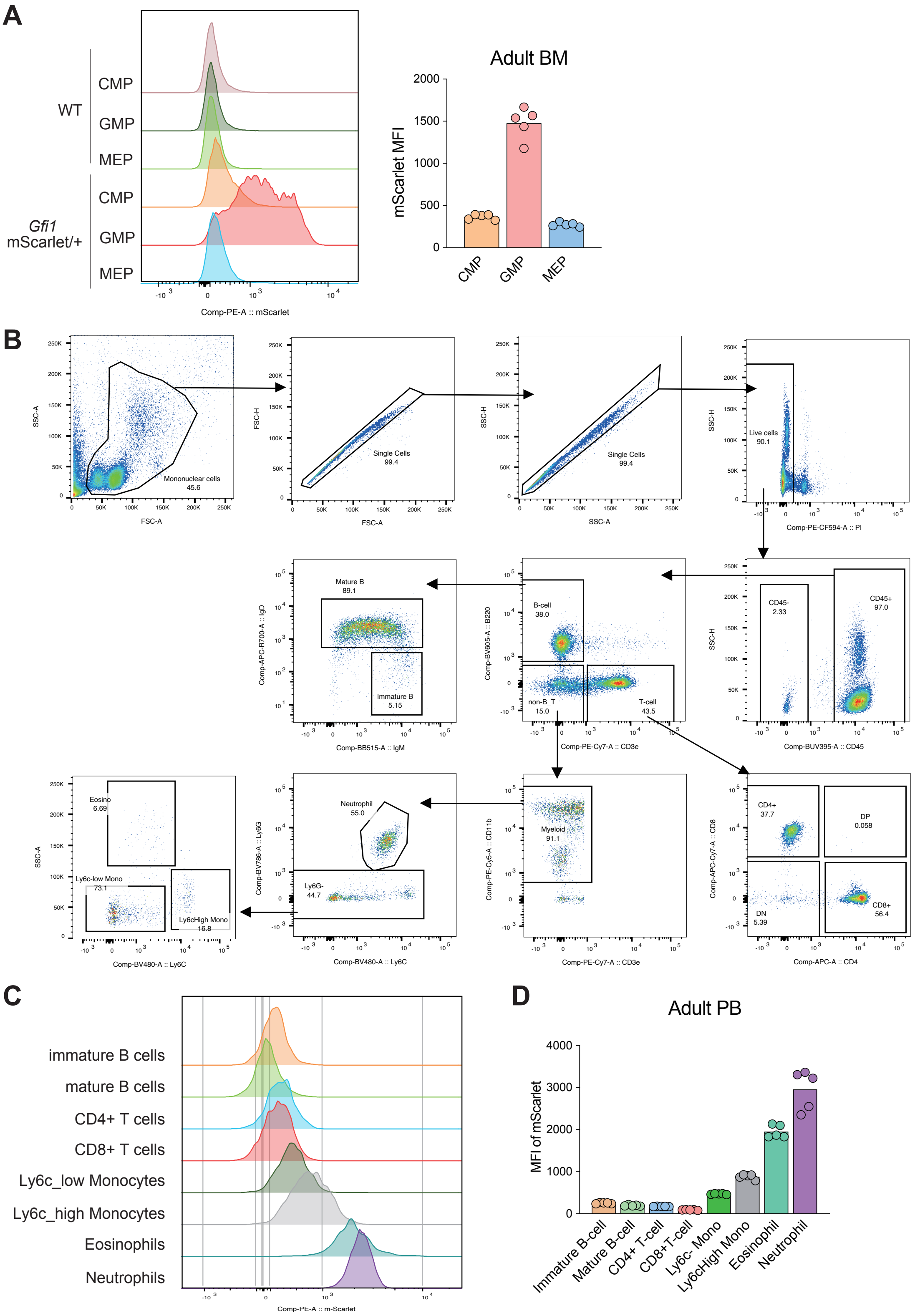
A
WT
CMP
GMP
MEP
Gfi1
mScarlet/+
CMP
GMP
MEP
Comp-PE-A :: mScarlet
Adult BM
mScarlet MFI
2000
1500
1000
500
0
CMP
GMP
MEP
B
Mononuclear cells
45.6
SSC-A
FSC-A
Single Cells
99.4
FSC-H
FSC-A
Single Cells
99.4
SSC-H
SSC-A
Live cells
90.1
SSC-H
Comp-PE-CF594-A :: PI
CD45-
2.33
CD45+
97.0
Comp-BUV395-A :: CD45
B-cell
38.0
non-B_T
15.0
T-cell
43.5
Comp-BV605-A :: B220
Comp-PE-Cy7-A :: CD3e
Mature B
89.1
Immature B
5.15
Comp-APC-R700-A :: IgD
Comp-BB515-A :: IgM
CD4+
37.7
DP
0.058
DN
5.39
CD8+
56.4
Comp-APC-Cy7-A :: CD8
Comp-APC-A :: CD4
Myeloid
91.1
Comp-PE-Cy5-A :: CD11b
Comp-PE-Cy7-A :: CD3e
Neutrophil
55.0
Ly6G-
44.7
Comp-BV786-A :: Ly6G
Comp-BV480-A :: Ly6C
Eosino
6.69
Ly6c-low Mono
73.1
Ly6cHigh Mono
16.8
SSC-A
Comp-BV480-A :: Ly6C
C
immature B cells
mature B cells
CD4+ T cells
CD8+ T cells
Ly6c_low Monocytes
Ly6c_high Monocytes
Eosinophils
Neutrophils
Comp-PE-A :: m-Scarlet
D
Adult PB
MFI of mScarlet
4000
3000
2000
1000
0
Immature B-cell
Mature B-cell
CD4+ T-cell
CD8+T-cell
Ly6c- Mono
Ly6cHigh Mono
Eosinophil
Neutrophil

**Figure S10. Granulocyte-monocyte progenitors, neutrophils and eosinophils exhibit high *Gfi1* expression.**

**(A)** Adult bone marrow myeloid progenitor subsets (CMP, GMP, MEP) compared between WT and *Gfi1*-mScarlet mice. Histograms and MFI analyses demonstrate reporter signals showing differences in *Gfi1* expression within the adult LK compartment. **(B)** Flow cytometric gating strategy of adult peripheral blood, identifying each labeled population in its respective gate. **(C-D)** Representative histograms showing Gfi1-mScarlet reporter fluorescence in peripheral blood subfractions of Gfi1-mScarlet mice (C) and mean fluorescence intensity (MFI) quantification for each population (D).

### Figure S11.

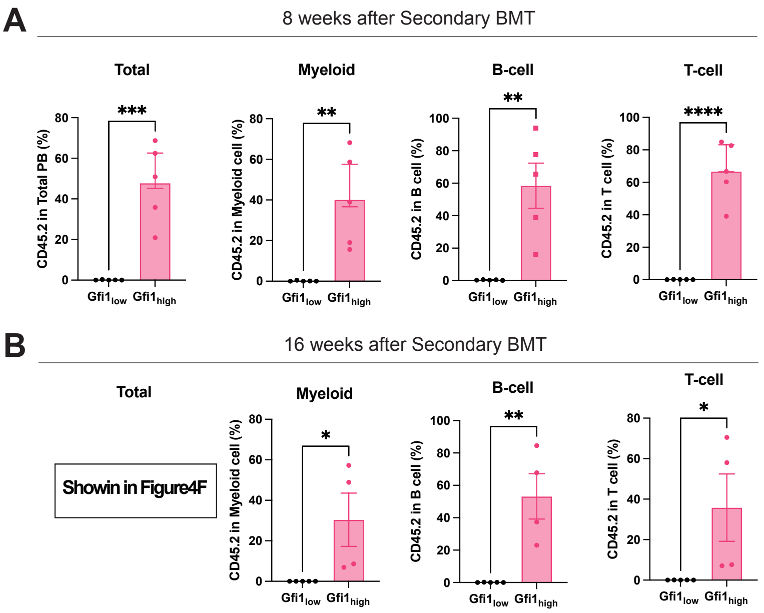


**Figure S11. Secondary transplantation analysis of adult BM HSCs.**

**(A-B)** Peripheral blood chimerism at 2 months **(A)** and 4 months **(B)** after secondary transplantation of adult BM *Gfi1*-high and *Gfi1*-low HSCs. Chimerism is shown for total cells, myeloid cells, B cells, and T cells.

### Figure S12.

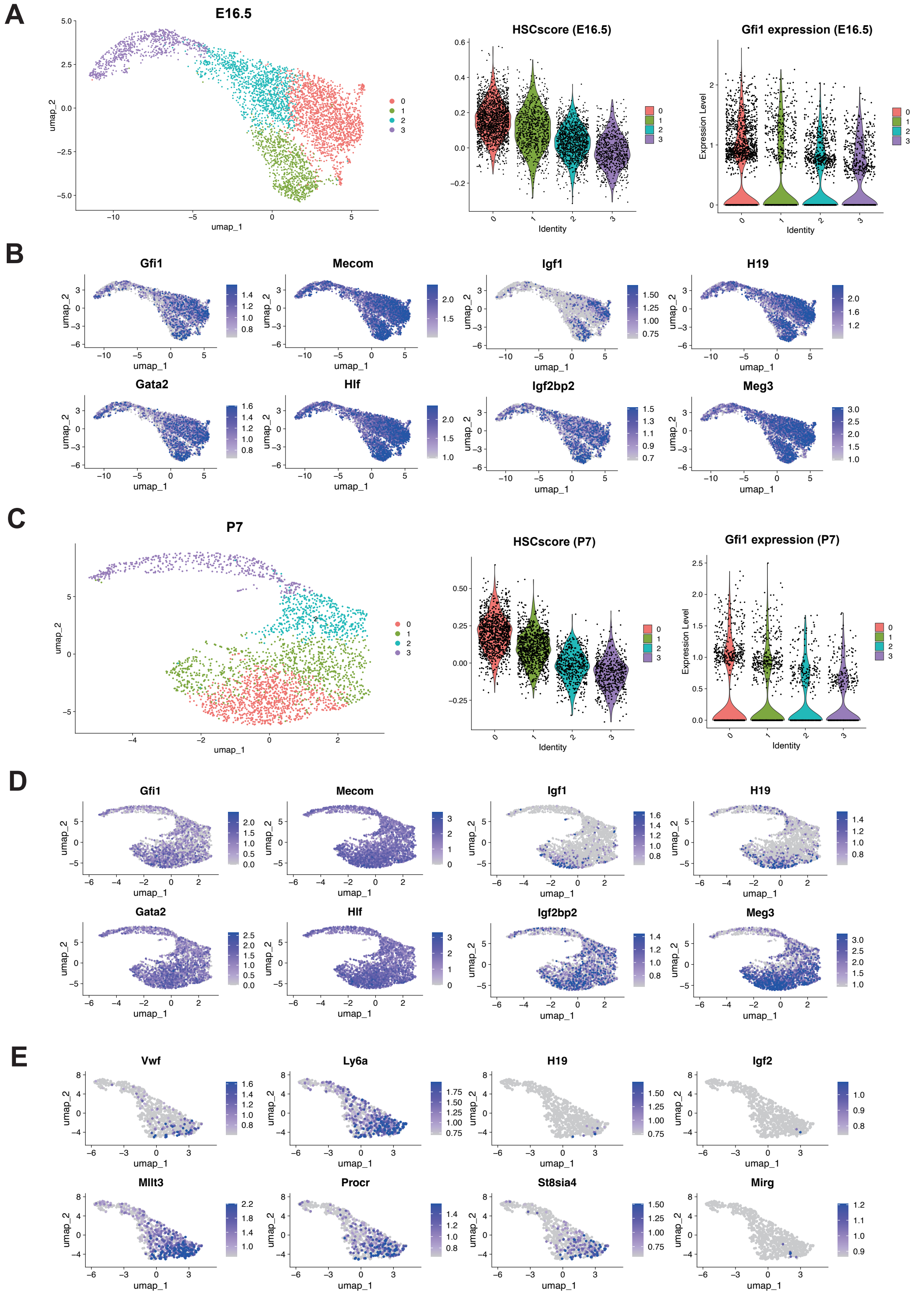
A
E16.5
HSCscore (E16.5)
Gfi1 expression (E16.5)
B
Gfi1
Mecom
Igf1
H19
Gata2
Hlf
Igf2bp2
Meg3
C
P7
HSCscore (P7)
Gfi1 expression (P7)
D
Gfi1
Mecom
Igf1
H19
Gata2
Hlf
Igf2bp2
Meg3
E
Vwf
Ly6a
H19
Igf2
Mllt3
Procr
St8sia4
Mirg

**Figure S12. Single-cell transcriptomic landscape of Gfi1-associated heterogeneity throughout HSC development.**

**(A)** UMAP visualization of fetal liver (E16.5) HSCs extracted from an integrated public scRNA-seq dataset[14]. Shown are the clusters identified by unsupervised re-clustering of the E16.5 HSC subset (left), dormant HSC scores (MolO scores[16]) across clusters (middle), and the distribution of Gfi1 expression in those clusters (right).

**(B)** Feature plots on the UMAP in (A) showing the expression patterns of HSC-associated transcription factors (*Gfi1*, *Mecom*, *Gata2*, and *Hlf*), alongside FL-enriched genes (*H19*, *Igf1*, *Meg3*, and *Igf2bp2*) across E16.5 HSC clusters. **(C)** UMAP representation of early postnatal (P7) HSCs analyzed as in (A), with dormant HSC scores (middle) and *Gfi1* expression patterns (right) **(D)** Feature plots on the UMAP in (C) showing the same HSC-associated transcription factors and FL-enriched genes examined in (B). **(E)** Feature plots of adult bone marrow HSCs showing HSC-associated markers (Vwf, Ly6a, Mllt3, and Procr) and FL-associated genes (*H19*, *Igf2*, *St8sia4*, and *Mirg*) whose expression is selectively enriched within *Gfi1*-high HSC clusters.

#### Figure S13.

A

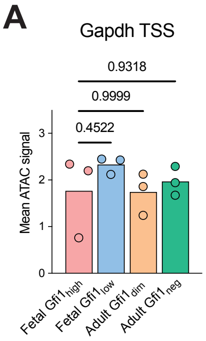


B

| Gene | Region | Coordinates |
|---|---|---|
| *Gfi1* | TSS | chr5:107,723,806–107,727,805 |
| | -33.5 | chr5:107,758,201–107,760,200 |
| | 26.5 | chr5:107,698,288–107,700,287 |
| *Gata2* | -83 | chr6:88,116,814-88,117,272 |
| *Gfi1b* | 16 | chr2:28,599,093–28,600,045 |
| *Gapdh* | TSS | chr6:125,164,458–125,168,457 |

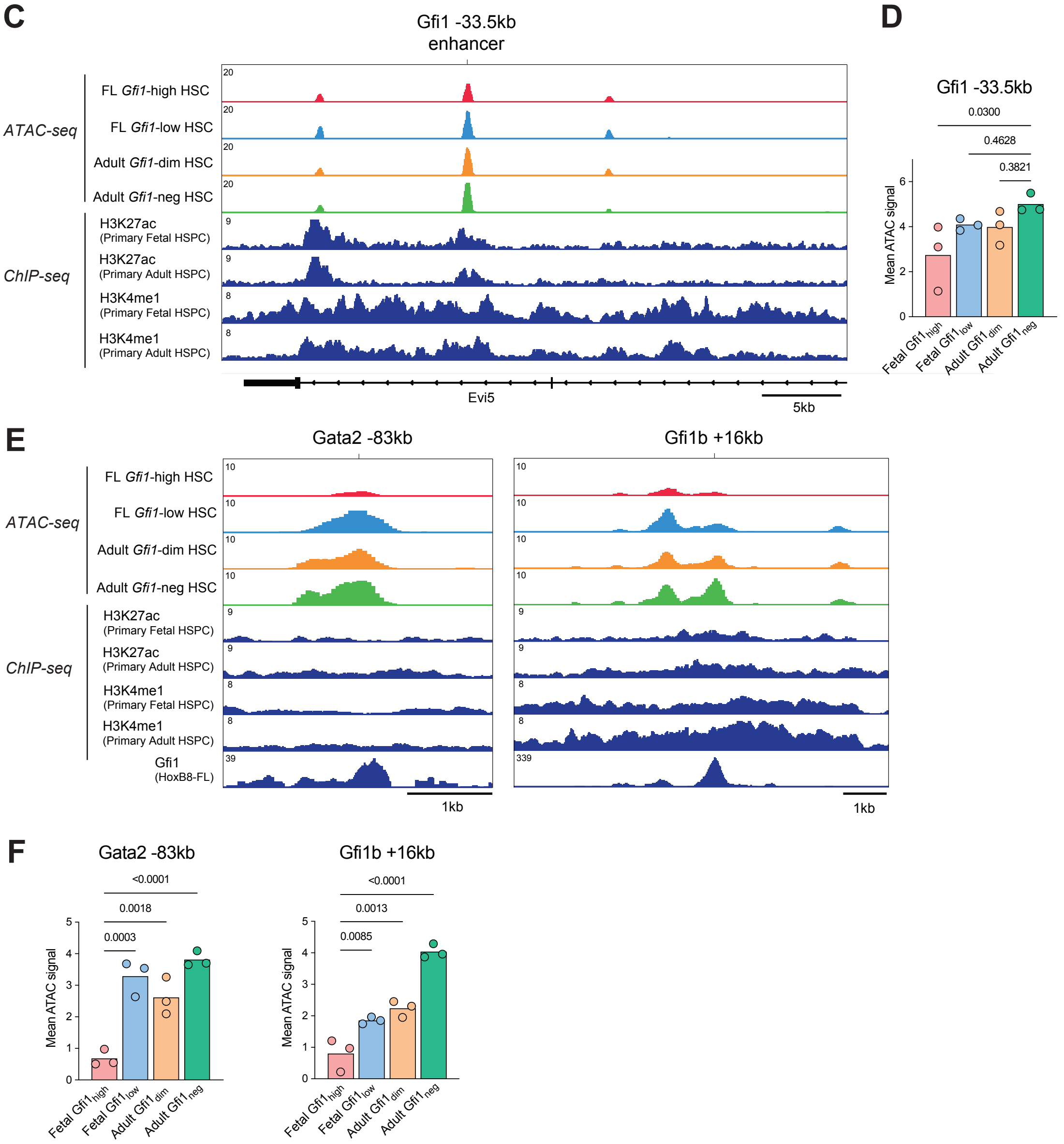

**Figure S13. Chromatin accessibility at selected regulatory regions was assessed in fetal and adult HSCs.**

ATAC-seq analyses were performed in *Gfi1*-high and *Gfi1*-low HSCs isolated from E19.5 FL and adult BM. **(A)** Quantification of mean CPM-normalized ATAC-seq signals at the *Gapdh* TSS (±2 kb). **(B)** Genomic regions used for ATAC-seq quantification, including gene names, region, and corresponding mm10 coordinates. **(C, D)** Mean CPM-normalized ATAC-seq tracks at the *Gfi1* -33.5 kb developmental hematopoietic enhancer, together with public H3K27ac and H3K4me1 ChIP-seq profiles from fetal and adult primary HSPCs (GSE128760[14]) **(C)**, and quantification of ATAC-seq signals within the ±1 kb region **(D)**. **(E, F)** Mean CPM-normalized ATAC-seq tracks at the *Gata2* −83 kb regulatory element and *Gfi1b* +16 kb enhancer, together with public H3K27ac and H3K4me1 ChIP-seq profiles from fetal and adult HSPCs (GSE128760[14]) and Gfi1 ChIP-seq profiles from Hoxb8-FL cells (GSE84328[21]) **(E)**, and quantification of ATAC-seq signals at both regions **(F)**.
